# Cetylpyridinium chloride (CPC) reduces zebrafish mortality from influenza infection: Super-resolution microscopy reveals CPC interference with multiple protein interactions with phosphatidylinositol 4,5-bisphosphate in immune function

**DOI:** 10.1101/2021.10.08.463687

**Authors:** Prakash Raut, Sasha R. Weller, Bright Obeng, Brandy L. Soos, Bailey E. West, Christian M. Potts, Suraj Sangroula, Marissa S. Kinney, John E. Burnell, Benjamin L. King, Julie A. Gosse, Samuel T. Hess

## Abstract

The COVID-19 pandemic raises significance for a potential influenza therapeutic compound, cetylpyridinium chloride (CPC), which has been extensively used in personal care products as a positively-charged quaternary ammonium antibacterial agent. CPC is currently in clinical trials to assess its effects on severe acute respiratory syndrome coronavirus 2 (SARS-CoV-2) morbidity. Two published studies have provided mouse and human data indicating that CPC may alleviate influenza infection, and here we show that CPC (0.1 μM, 1 hour) reduces zebrafish mortality and viral load following influenza infection. However, CPC mechanisms of action upon viral-host cell interaction are currently unknown. We have utilized super-resolution fluorescence photoactivation localization microscopy to probe the mode of CPC action. Reduction in density of influenza viral protein hemagglutinin (HA) clusters is known to reduce influenza infectivity: here, we show that CPC (at non-cytotoxic doses, 5-10 µM) reduces HA density and number of HA molecules per cluster within the plasma membrane of NIH-3T3 mouse fibroblasts. HA is known to colocalize with the negatively-charged mammalian lipid phosphatidylinositol 4,5-bisphosphate (PIP_2_); here, we show that nanoscale co-localization of HA with the PIP_2_-binding Pleckstrin homology (PH) reporter in the plasma membrane is diminished by CPC. CPC also dramatically displaces the PIP_2_-binding protein myristoylated alanine-rich C-kinase substrate (MARCKS) from the plasma membrane of rat RBL-2H3 mast cells; this disruption of PIP_2_ is correlated with inhibition of mast cell degranulation. Together, these findings offer a PIP_2_-focused mechanism underlying CPC disruption of influenza and suggest potential pharmacological use of this drug as an influenza therapeutic to reduce global deaths from viral disease.

## Introduction

Recent intriguing studies (Mukherjee *et al.*, 2017; Popkin *et al.*, 2017) suggest that the classic antibacterial agent cetylpyridinium chloride (CPC) may fight influenza infections in mice and humans. However, CPC antiviral mechanisms of action in any mammalian host are completely unknown (Pubmed Sept. 2021).

CPC is a positively-charged, quaternary ammonium (quat) antimicrobial agent used extensively in cleaning and personal care products, including moisturizers, cleansing wipes, mouthwashes, and various types of toothpaste (Mao *et al.*, 2020) at concentrations of 1500-3000 μM (Rawlinson *et al.*, 2008). Widespread use of CPC is mostly attributed to its antibacterial action, for example against plaque and gingivitis (Witt *et al.*, 2005) via oral care products. While very little information has been published regarding exposure levels and metabolism of CPC in humans or other eukaryotes following product exposure, CPC is retained in the oral mucosa for long periods of time after product usage, as indicated by slow release of substantial CPC concentrations into the saliva (Bonesvoll and Gjermo, 1978). Alternative therapeutic uses for this over-the-counter drug, beyond antibacterial action, could be powerful.

As a quat, CPC consists of a monocationic head group at the end of a hydrophobic chain. To impede bacterial infection, CPC can act as a detergent to solubilize individual bacterial cells, causing release of cytoplasmic contents and bacterial cell death (Gilbert and Moore, 2005). This detergent action is realized when CPC is present *above* its critical micelle concentration (CMC) of ∼900 µM in pure water (Mukerjee and Mysels, 1971), such as at concentrations found in common consumer products (1500-3000 µM). Different types of salt buffers can significantly lower the CMC (Abezgauz *et al.*, 2010), but numerous studies which determined the CMC by various distinct methods and with a wide range of buffer components and temperatures all found a CMC of CPC to be ∼600-900 µM (Mandal and Nair, 2002; Varade *et al.*, 2005; Abezgauz *et al.*, 2010; Shi *et al.*, 2011). For example, around the CMC, CPC interferes with BODIPY-TR-cadaverine binding to gram-negative bacterial lipopolysaccharides (LPS) and to the gram-positive bacterial lipid lipoteichoic acid (LTA), suggesting that CPC may target these bacterial outer membrane lipids (Haught *et al.*, 2016). The compact, conical molecular shape of CPC and its hydrocarbon tail, which is similar in length to the tails of phosphatidylcholine, are features that likely contribute to its ability to insert itself into membranes (Arrigler *et al.*, 2005).

Below its CMC, CPC may also exert other types of antibacterial actions. For example, CPC concentrations ≥ 2 μM interfere with the LPS/LTA, as measured by the Limulus Amebocyte Lysate method (Haught *et al.*, 2016). Another indication of CPC binding to LPS is its ability, at concentrations as low as 16.5 µM, to inhibit LPS binding to Toll-like Receptor 4 on the surface of human cells (Haught *et al.*, 2016). As a quat, CPC may displace and replace divalent cations which normally stabilize the lipopolysaccharides within the gram–negative bacterial membrane and the negative charge of all bacterial cells’ surfaces (Vaara, 1992; Gilbert and Moore, 2005; Haught *et al.*, 2016; Mao *et al.*, 2020). In turn, CPC at concentrations of ∼ 24,000 µM hinders microbial membrane lipid integrity by electrostatic attraction and charge neutralization between oppositely charged components of the microbe and CPC resulting in aggregate formation (Yegin *et al.*, 2019). Therefore, metabolic and biosynthetic processes within bacteria may be perturbed at concentrations both lower and higher than the CMC. Thus, there are both alternative mechanisms of antibacterial action (vs. simple detergent solubilization above the CMC) and evidence of CPC binding to specific bacterial lipids. While cholesterol protects cell membranes from solubilization by quats, leading to greater resistance to quats in eukaryotic cells compared to bacterial cells (Watanabe and Regen, 1994; Marcotte *et al.*, 2005), there is otherwise a dearth of published literature regarding information on CPC interactions with eukaryotic cell lipids.

In addition to its antibacterial action, CPC also has a demonstrated potential as an antiviral agent (Popkin *et al.*, 2017; Seo *et al.*, 2019; Baker *et al.*, 2020; Koch-Heier *et al.*, 2021). To date, most studies exploring antiviral activity of CPC either have not included mechanistic information or have focused on direct interactions between CPC and the viral particle. To our knowledge, information on CPC effects on virus interaction with eukaryotic host cells is lacking. CPC was shown to be an effective inhibitor of Hepatitis B (HBV). The direct mechanism behind CPC’s HBV inhibition ability involves its prevention of capsid assembly via interaction with the viral nucleocapsid protein, which ultimately leads to reduced HBV biogenesis (Seo *et al.*, 2019). At 1500-2100 µM, CPC destroys SARS-CoV-2 viruses within tens of seconds (Koch-Heier *et al.*, 2021). CPC is believed to inhibit enveloped viruses in general by disrupting viral membranes (Baker *et al.*, 2020). In particular, CPC possesses virucidal activity against several strains of influenza virus, even at doses below its CMC (∼15-60 μM, with exposure times of a few minutes), apparently by disrupting the morphology of the viral envelope (Popkin *et al.*, 2017). The authors suggested that CPC may target the mammalian host cell-derived lipids within the viral envelope, such as phosphatidylethanolamine.

Furthermore, CPC administered orally to mice ameliorated influenza infection and reduced morbidity in mice (Popkin *et al.*, 2017). Annually, in the U.S. alone, there are typically 12,000 – 61,000 mortalities due to influenza (CDC, 2020). Thus, there is a current global need for novel forms of drug treatment (Antara, 2018) in order to combat annual seasonal cases of influenza, including those refractory to the annual vaccine. Intriguingly, CPC was tested in a human clinical trial to assess its effectiveness against upper respiratory infections (Mukherjee *et al.*, 2017). Very recently, CPC has been the subject of high-profile news reports for its use in multiple clinical trials against SARS-CoV-2. For example, one human clinical trial assessed use of CPC to reduce oral load of CPC in patients preparing to undergo dental procedures; the researchers found that CPC mouthwash decreased levels of SARS-CoV-2 in patients’ saliva following a 5-minute rinse, as compared to a 5-minute water rinse (Seneviratne *et al.*, 2021). Thus far, studies have indicated both that CPC may directly destroy viruses and that it may be an effective pharmacological antiviral within animals and humans.

However, the question remains--does CPC act as an antiviral by disrupting eukaryotic host cell/virus interactions? At low micromolar doses far below the CMC, CPC inhibited fusion between the SARS-CoV-2 viral membrane and HEK-293T human kidney cells (Muñoz-Basagoiti *et al.*, 2020). Another recent study showed that CPC, at concentrations of a few tens of micromolar or less, inhibits herpes simplex virus replication in human cells via CPC effects on the NF-κB pathway (Alvarez *et al.*, 2020). To our knowledge, these are the first reports of CPC disruption of a virus-host cell interaction (Green *et al.*, 2020).

There are no published studies regarding CPC effects on immune cell function (Pubmed); in the current study, we explore the effect of CPC on the function of mast cells. Mast cell signaling and function hinge upon the key role of phosphatidylinositol 4,5 bisphosphate (PIP_2_), a plasma membrane signaling lipid. Mast cells, found throughout human tissues, defend the body against bacterial (Johnzon *et al.*, 2016), viral (Dawicki and Marshall, 2007), and parasitic (Metcalfe *et al.*, 1997) infections as well as playing key roles in allergy, asthma (Galli *et al.*, 2005), and neurological function (Theoharides *et al.*, 2016). Mast cells undergo an allergen/antigen (Ag)-stimulated process called degranulation, in which they release bioactive substances including histamine and serotonin. Mast cell degranulation begins when a multivalent Ag binds to and mediates cross-linking of IgE-bound FcεRI receptors leading to a phosphorylation cascade in which spleen tyrosine kinase (Syk) and phospholipase C gamma (PLCγ) are activated respectively (Kinet, 1999). The activated PLCγ cleaves phosphatidylinositol 4,5 bisphosphate (PIP_2_) to generate diacylglycerol (DAG) and inositol 1,4,5-trisphosphate (IP_3_). The IP_3_ binds to its receptor on the endoplasmic reticulum (ER) membrane, triggering an efflux of Ca^2+^ from the ER internal stores into the cytosol (Berridge, 1993). The subsequent activated influx of Ca^2+^ into the cytosol is termed store-operated calcium entry (SOCE) (Putney, 1986). SOCE is a core mediator of the mast cell degranulation pathway and necessary for downstream events such as protein kinase C (PKC) activation (Ozawa *et al.*, 1993), that lead to degranulation. PIP_2_-dependent SOCE is also key for T Cell function (Punt *et al.*, 2019; Trebak and Kinet, 2019). In this study, we examine the effects of CPC on degranulation, using the RBL-2H3 mast cell type, which react to environmental stimuli in a fashion analogous to the biochemical responses of primary bone marrow-derived mast cells (Zaitsu *et al.*, 2007; Thrasher *et al.*, 2013; Alsaleh *et al.*, 2016). In this study, we examine CPC effects on degranulation to test our hypothesis that CPC interferes with PIP_2_-dependent cellular processes.

PIP_2_ is a negatively-charged minority lipid residing in the inner leaflet of the mammalian plasma membrane (PM) (McLaughlin and Murray, 2005). In addition to immune cell signaling, PIP_2_ is also involved in viral infectivity (Mucksch *et al.*, 2019). PIP_2_ can modify the orientation of nearby phosphatidylcholine lipids within a bilayer (Poyry and Vattulainen, 2016). PIP_2_ can electrostatically bind to the myristoylated alanine-rich C-kinase substrate (MARCKS) within the inner leaflet of the plasma membrane (PM) (Heo *et al.*, 2006). MARCKS is a substrate for enzyme PKC, which phosphorylates MARCKS, causing MARCKS dissociation from the PM in an oscillatory pattern to the cytosol (Gadi *et al.*, 2011). The MARCKS construct used in this study, called monomeric red fluorescent protein (mRFP)-MARCKS-ED, is composed of basic amino acids within its effector domain (ED) which bind electrostatically to PIP_2_ at the PM (Gadi *et al.*, 2011) and which can also bind equally strongly to PI(3,4)P_2_ (Wang *et al.*, 2001). Additionally, PIP_2_ electrostatically binds to many other proteins, including those with Pleckstrin Homology (PH) domains, via conserved, positively-charged lysine or arginine residues, which bind to the 4- and 5-monoesters of PIP_2_ specifically and more tightly than to other phosphorylated inositols (Harlan *et al.*, 1995). In this study, a fluorescent protein-tagged PH domain construct, PAmKate-PH, is used to probe CPC effects on PIP_2_-binding proteins (Curthoys *et al.*, 2019).

A central hypothesis of this study is that, via its positive charge, CPC biochemically targets the negatively-charged eukaryotic lipid PIP_2_ and, therefore, interferes with PIP_2_-protein interactions and function. Evidence in the literature points to a role for positively-charged cations and even antibiotics including neomycin in electrostatically blocking interactions between PIP_2_ and PIP_2_-binding proteins (Suh and Hille, 2008; Seo *et al.*, 2015). The clustering of PIP_2_ can have many functional ramifications, including modulation of a variety of proteins relevant to cell signaling (Hammond, 2016). PIP_2_ clustering can modulate PM curvature in conjunction with cations such as Ca^2+^, which can in turn disrupt cellular processes (Graber *et al.*, 2017) such as degranulation (Kazama *et al.*, 2013). Formation of PIP_2_ clusters can be promoted by Ca^2+^ and other multivalent metal ions (Wen *et al.*, 2018) and by detergents like Triton X-100 (van Rheenen *et al.*, 2005).

The influenza viral protein hemagglutinin (HA), which catalyzes viral fusion and entry into the mammalian host cell, has been shown via super-resolution imaging to colocalize with PIP_2_ on the nanoscale, and HA and PIP_2_ clustering are interdependent (Curthoys *et al.*, 2019), mutually altering cluster area and density (Curthoys *et al.*, 2019). Importantly, HA must cluster within the mammalian host cell PM for assembly, HA clusters are needed for fusion competence of progeny viruses, and smaller or less-dense HA clusters are correlated with lower influenza infectivity (Ellens *et al.*, 1990; Takeda *et al.*, 2003). We hypothesize that PIP_2_ interacts with HA through the HA cytoplasmic domain, which is composed of 10-11 amino acids with a net charge of +2 at physiological pH, and which contains three highly-conserved acylated cysteines adjacent to its two positively-charged amino acids (Simpson and Lamb, 1992; Veit *et al.*, 2013; Curthoys *et al.*, 2019). PIP_2_ may also interact indirectly with HA via interactions between the PM, cortical actin cytoskeleton, and other membrane-associated proteins (Raucher *et al.*, 2000; Curthoys *et al.*, 2019). Specifically, HA can colocalize with actin-binding proteins (Guerriero *et al.*, 2006). Thus, these findings suggest that PIP_2_ plays a regulatory role in HA clustering and function and thus in influenza infectivity.

Super-resolution imaging was necessary for the work in this study because PIP_2_ and HA clusters are typically too small to image using diffraction-limited microscopy (Curthoys *et al.*, 2019). Fluorescence photoactivation localization microscopy (FPALM; (Hess *et al.*, 2006) was used in order to visualize these molecular interactions on the nanoscale. FPALM can detect molecular distributions in cells and other biological systems at a resolution of ∼10-20 nm (Hess *et al.*, 2006; Hess *et al.*, 2009). FPALM uses repeated cycles of activation, localization, and photobleaching in conjunction with high sensitivity fluorescence imaging, to identify and localize numerous molecules within a sample (Hess *et al.*, 2006), and is compatible with live-cell (Hess *et al.*, 2007), 3-dimensional (Juette *et al.*, 2008) and multicolor (Gunewardene *et al.*, 2011).

In this study, we hypothesize that positively-charged CPC biochemically targets the negatively-charged eukaryotic lipid PIP_2_ and, therefore, interferes with PIP_2_-protein interactions and function at ∼1,000-fold lower concentrations than those found in personal care products. In order to address these questions, this study will assess CPC effects on the nanoscale distribution, colocalization, clustering, and density of three PIP_2_-binding proteins (MARCKS, HA, and PH) in the PM of eukaryotic cells. Consequent to CPC disruption of these PIP_2_-protein interactions, functional effects on mast cells and to influenza infection in zebrafish are also examined. In this study, we are promulgating a mechanism by which CPC affects the mammalian cell itself, to reduce influenza morbidity. We are also extending the earlier finding of CPC reduction of influenza morbidity in mice (Popkin *et al.*, 2017) to the zebrafish model system, which has been used previously to study influenza infection (Gabor *et al.*, 2014). We hypothesize that CPC is not simply a virucidal agent; rather, at low- and sub -micromolar doses, CPC is affecting the host cell-virus interaction by interfering with the phosphoinositide PIP_2_ and with PIP_2_-HA interactions. These findings reveal CPC as a potential therapeutic to treat PIP_2_-related diseases like influenza and also may modulate other PIP_2_-dependent eukaryotic cell functions.

## Methods

### Chemical and Reagents

Cetylpyridinium chloride (CPC; 99% purity, VWR; CAS no. 123-03-5) was prepared at 150 µM in a pre-warmed Tyrodes buffer (recipe given in (Hutchinson *et al.*, 2011)) and vortexed. Next, the preparation was sonicated (Branson 1200 ultrasonic cleaner; Branson Ultrasonics, Danbury, CT, USA) at 37 °C for 20 minutes, protected from light, then vortexed again. The solution was then poured into a sterile Erlenmeyer flask for continual stirring until usage, protected from light. A control Tyrodes buffer solution was prepared in tandem. Following dilution in pre-warmed Tyrodes buffer, exact concentrations were determined using UV-Vis spectrophotometry (Weatherly *et al.*, 2013) and the Beer-Lambert equation (A_260_ =ε_260_ℓc), using an ε_260_ of 4389 M^-1^ cm^-1^ (Bernauer *et al.*, 2015). Bovine serum albumin (BSA) was added to create a final solution of CPC in BT (Tyrodes buffer containing BSA (Hutchinson *et al.*, 2011)). For all cell culture experiments, CPC was administered via BT. For zebrafish experiments, CPC was administered via egg water (60 μg/ml Instant Ocean sea salts in autoclaved Milli-Q water, pH 7.0; Aquarium Systems, Mentor, OH) (with no BSA), and sonication and UV-Vis were performed similarly as above. Fresh CPC stocks were prepared on each experimental day.

### Cell Culture

#### RBL-2H3 Cell Culture

RBL-2H3 mast cells were cultured as described (Hutchinson *et al.*, 2011).

#### NIH-3T3 Cell Culture

NIH-3T3 mouse fibroblast cells (ATCC, CRL-1658) were cultured as described (Curthoys *et al.*, 2019).

#### MDCK London Cell Culture

MDCK London cells (passage 3) were grown at 37 °C with 5% CO_2_ in T-175 flasks in minimal essential medium (MEM), containing final percentages/concentrations of the following: 5% newborn calf serum (NCS), 2% heat-inactivated fetal bovine serum (FBS), 0.23% sodium bicarbonate solution, 2% MEM amino acids (from 50X stock), 1% MEM vitamins (from 100X stock), 4 mM L-glutamine, and 1% Antibiotic-Antimycotic (from 100X stock). All media and reagents for MDCK culture were from ThermoFisher Scientific unless otherwise noted. Cells were maintained through washing twice with 1x Dulbecco’s phosphate-buffered saline (dPBS, pH 7.4) to remove traces of NCS and FBS, trypsinizing with 0.25% trypsin-EDTA with phenol red, and passaging in a 1:6 dilution every 2-3 days. Virus-infected cells were grown in MEM-BSA/TPCK: this media is similar to the MEM media described above but supplemented with Tosyl phenylalanyl chloromethyl ketone (TPCK) trypsin (Worthington Chemical Corporation) and with Bovine Albumin Fraction V (7.5% solution) instead of NCS and FBS.

### MARCKS Assay in RBL-2H3 Mast Cells

RBL-2H3 mast cells were transiently transfected with monomeric red fluorescent protein (mRFP)-MARCKS-ED plasmid (a gift from Dr. Barbara Baird and Dr. David Holowka, Cornell University; “MARCKS”, (Gadi *et al.*, 2011)) via electroporation using Amaxa Nucleofactor kit T (Lonza) as described in (Weatherly *et al.*, 2018). After electroporation, cells were plated at 1.7 x 10^5^ cells per well (in an eight-well ibidi plate) in a phenol red-free media and incubated overnight at 5% CO_2_/ 37 ℃. The next day, the spent media was removed, and cells were incubated with 200 µl of 0, 5, or 10 µM CPC in BT for 30 minutes. Following this incubation, cells were washed in BT, 200 µl BT was added to each well, and images were taken immediately using confocal microscopy. See “Confocal Microscopy” below for imaging details.

### Confocal Microscopy

For RBL-2H3 cells transiently transfected with monomeric red fluorescent protein (mRFP)-MARCKS-ED plasmid, an Olympus FV-1000 confocal microscope with an Olympus IX-81 inverted microscope, with a 1-mW HeNe-Green laser (543 nm excitation and 560-660 nm emission filter) were used to collect images. An oil immersion 100 x objective with NA 1.4 and image acquisition speed of 2 μs/pixel were used to collect the images. Using the ibidi plate heating system, imaging was conducted at 37 ℃.

### Automated Image Analysis

Fiji ImageJ software (NIH) was used to analyze confocal microscopy images of RBL-2H3 cells transfected with monomeric red fluorescent protein (mRFP)-MARCKS-ED. Images were converted to 8-bit images, channels split, and further analysis done using the fluorescent channels. Pseudo flatfield correction was used to subtract background, appropriate threshold was applied, and binary masks generated for both the whole cell and the cytoplasm of a given transfected cell. The masks of the whole cell and of the cytoplasm were applied to a given transfected cell in the fluorescent channel to generate regions of interest (ROI) for the whole cell and the cytoplasm, respectively. Next, the area, integrated density, and mean fluorescence per pixel of the whole cell and the cytoplasm were measured from the ROI. The mean fluorescence intensity per pixel of the PM values were generated by 1.) subtracting the integrated density of the cytoplasm from the integrated density of the whole cell, 2.) subtracting the area of the cytoplasm from the area of the whole cell, and 3.) dividing the result of 1.) by the result of 2.). The PM/cytoplasm mean fluorescence per pixel were calculated by dividing the mean fluorescence of the PM by the mean fluorescence of cytoplasm, and results analyzed in GraphPad Prism.

### Degranulation Assay

Degranulation response was measured in RBL-2H3 mast cells as in (Weatherly *et al.*, 2013). This assay indicates the level of degranulation occurring by measuring the release of enzyme β-hexosaminidase into the surrounding environment. The β-hexosaminidase substrate utilized for detection was 4-methyllumbelliferyl-*N*-acetyl-β-D-glucosaminide (4-MU), a synthetic substrate that releases a fluorescent product upon enzymatic cleavage. We validated that there is no CPC effect on β-hexosaminidase’s ability to bind to its substrate, 4-MU (Figure S1).

Treatment groups included IgE-sensitized, antigen (DNP-BSA; Ag)-stimulated treatment cells, to be exposed to CPC dilutions; spontaneous cells, an indicator of the basal amount of degranulation that normally occurs; high-control cells, cells exposed to Triton-X 100 detergent to lyse all cells and maximize potential β-hexosaminidase release; and no-cell background wells, wells that received no cells that can be used to subtract out background fluorescence. The data were analyzed by subtracting the background fluorescence from all readings. The resultant values were then divided by the average high-control fluorescence values to obtain a “percentage degranulation” value. Individual experiments are normalized to the 0 μM CPC dose before averaging multiple days of experiments. Each treatment group was performed in experimental triplicate, with three wells of cells per treatment per experiment.

### CPC Cytotoxicity and Survivability on RBL-2H3 cells

Trypan blue exclusion and lactate dehydrogenase (LDH) cytotoxicity assays were used to assess cytotoxicity and survivability on RBL-2H3 cells exposed to CPC. The trypan blue exclusion assay was performed exactly as in Hutchinson 2011 (Hutchinson *et al.*, 2011), except with overnight plating of 1.2 x 10^6^ cells per well of six-well Grenier CELLSTAR plates and with CPC exposure instead of arsenic.

Cytotoxicity was also assessed based upon LDH release from cells (into the supernatant), using an absorbance-based cytotoxicity detection kit per instructions (Sigma), as described in Hutchinson 2011 (Hutchinson *et al.*, 2011). We confirmed that CPC does not interfere with the LDH assay (Figure S2).

### Fluorescence Photoactivation Localization Microscopy (FPALM) Imaging and Processing

For sample preparation, cells were seeded in growth media (DMEM with Glucose without phenol red, Lonza) with 10% Calf Bovine Serum (ATCC) and antibiotics (Penicillin Streptomycin, 100 ug/ml) at 35,000 cells/ml overnight in 35 mm petri dishes (MatTek). After 20-24 hours, cells were transfected using Lipofectamine 3000 (Invitrogen) with 2 µg of total DNA (1µg of HA-Dendra2 (Gudheti *et al.*, 2013) and 1µg of PAmKate-PH (PLC-δ) (Curthoys *et al.*, 2019)) and treated with Control (0 µm CPC in BT vehicle), 5µM CPC in BT, or 10 µM CPC in BT for 1 hour in the incubator under same conditions. After one hour, cells were washed twice with PBS (Sigma-Aldrich), fixed at room temperature in 4% paraformaldehyde (PFA) (Alfa Aesar) for 10-15 minutes, and then washed again with PBS two times.

Two color FPALM was performed by previously published methods (Hess *et al.*, 2006; Gunewardene *et al.*, 2011; Curthoys *et al.*, 2019; Sangroula *et al.*, 2020). Briefly, lasers with wavelengths λ = 405 nm (CrystaLaser, 5mW) for the activation of Dendra2 molecule (Gurskaya et al., 2006) and λ = 558 nm for the readout (CrystaLaser, 100 mW) were focused in the back-aperture plane of an oil immersion objective (Olympus 60X 1.45NA) using a second lens (f = 350 mm, Thorlabs) at one focal length away from the back aperture. To match the polarization of the readout laser, the activation laser was passed through a half wave plate (Newport, 10RP42-1) and a linear polarizer (Newport, 5511). For better activation and readout, both lasers were then converted to elliptical (approximately circular) polarization by passing them through a quarter wave plate (Newport, 10RP54-1B). Laser power was recorded by the power meter (Thor labs) to be ∼13 mW for the readout laser and about ∼75 µW for the activation laser, respectively, after passing straight through the objective lens. The position of the beam in the objective back aperture was then translated to allow total internal reflection fluorescence (TIRF) imaging. Fluorescence emission was collected by the objective and filtered through a quad-band dichroic (Di01 R405/488/561/635-25×36), and by 405 nm and 561 nm notch filters (Semrock: NF03-561E-25). After the dichroic and notch filter, fluorescence emitted through the tube lens was magnified ∼2X using successive achromatic lenses with focal lengths +20 mm and +40 mm. Fluorescence then reached a second dichroic (Semrock, FF580-FDi02-t3) within the multi-color detection module, which reflected λ < 595 nm and transmitted λ > 595 nm, thus producing two wavelength ranges which were simultaneously imaged onto adjacent regions of the camera sensor. Fluorescence from the transmitted channel passed through an ET630/92 filter (Semrock, FF01-630/69-25) and fluorescence from the reflected channel passed through a 585/40 filter (Semrock, FF01-585/40-25) before reaching the camera (iXon+ DU897DCS-BV, Andor Scientific, Dublin, Ireland). Typically, ten thousand frames were recorded at ∼32 Hz and EM gain of 200.

### FPALM Data Analysis

Point spread functions recorded in the raw images were background subtracted (Sternberg, 1983) and localized by fitting into the 2D Gaussian function (Hess *et al.*, 2006). After localization, images were further analyzed for drift subtraction and bleed through correction (Kim *et al.*, 2013). Localizations were assigned to either of two channels according to their alpha values (Gunewardene *et al.*, 2011; Curthoys *et al.*, 2013) and further processed through custom built Matlab code for the quantitative analysis of clusters.

We used single linkage cluster analysis (SLCA) (Greenfield *et al.*, 2009; Gudheti *et al.*, 2013; Curthoys *et al.*, 2019; Sangroula *et al.*, 2020; Sneath, 1957) for cluster identification. This technique detects molecules within a maximum distance d_max_ of each other and assigns them to the same cluster. Clusters are analyzed further if they contain at least a minimum total number of molecules. For this analysis, d_max_ = 35 nm. To be considered for analysis, a threshold for the minimum number of molecules per cluster was required to be N_min_≥ 50.

The mean density of HA molecules and the mean density of PH molecules was obtained computationally. The process involves binning the molecules into a grid of squares 35 nm X 35 nm and counting the localizations of each species within each bin to create a density map. The density map was then convolved with a circle of 50 nm radius. A cell mask was calculated by automatically detecting the edge of the cell which involves dilating the convolved bins and filling the interior to approximately map the edge of the cell. After this cell area was computed and total number of localizations of each species was divided by the cell area to obtain mean density. Mean density was then averaged over all the cells and plotted as a function of CPC treatment.

The mean molecule number per grid pixel for HA within PH clusters (N_HA-PH_) and PH within HA clusters (N_PH-HA_) was determined from two-color FPALM results. This Mean Pixel Sum analysis (in PH channel within HA cluster or in HA channel within PH cluster) was performed by binning the localized molecules within a grid of 35 nm X 35 nm squares, after separation of their color according to their alpha values and after bleedthrough correction. Pixels (within the grid) were identified as containing a cluster for the first species (PH or HA) if they contained five or more molecules. After this identification, all the localizations of the second species (HA or PH) were summed over only those pixels that were previously identified to have a cluster of the first species, and this value (the summed number of localizations per pixel) was then averaged over all the cells. These resultant values were plotted as “mean pixel sum.”

### Zebrafish Care and Maintenance

AB Zebrafish (*Danio rerio*) used in the study were raised and housed following the recommendations in the Guide for the Care and Use of Laboratory Animals of the National Institutes of Health. The protocols used in this study were approved by the University of Maine Institutional Animal Care and Use Committee (IACUC) (Protocol Number A2018-01-01). Zebrafish were maintained in the Zebrafish Facility at the University of Maine. The Zebrafish Facility was maintained in accordance to IACUC standards. Embryos were collected from natural spawnings of adult AB zebrafish using varying sets of females and males. Fertilized eggs were collected as a pool from ∼20 females and 15 males per experiment. Embryos were raised in sterilized egg water at 33 °C.

### Influenza Type A Virus (IAV) and Survivability Curves

Influenza infection studies were performed as outlined in Figure 1. Purified Influenza A/Puerto Rico/8/34 H1N1 (APR8) virus was purchased from Charles River Laboratories (catalog number 10100374, Wilmington, MA), stored at −80 °C, thawed at room temperature, and diluted in cold sterile HBSS at a ratio of 87% virus to 13% diluent. Wild-type zebrafish embryos were injected into the duct of Cuvier (DC) as described in “Microinjection of IAV.” For survival studies, egg water was changed daily. All fish were monitored for morbidity, and moribund fish were euthanized with an overdose of sodium bicarbonate-buffered MS-222 (Tricaine S) solution (300 mg/L; Syndel US, Ferndale, WA). Mortality was recorded from 1-5 days post-infection (dpi).

**Figure 1.**
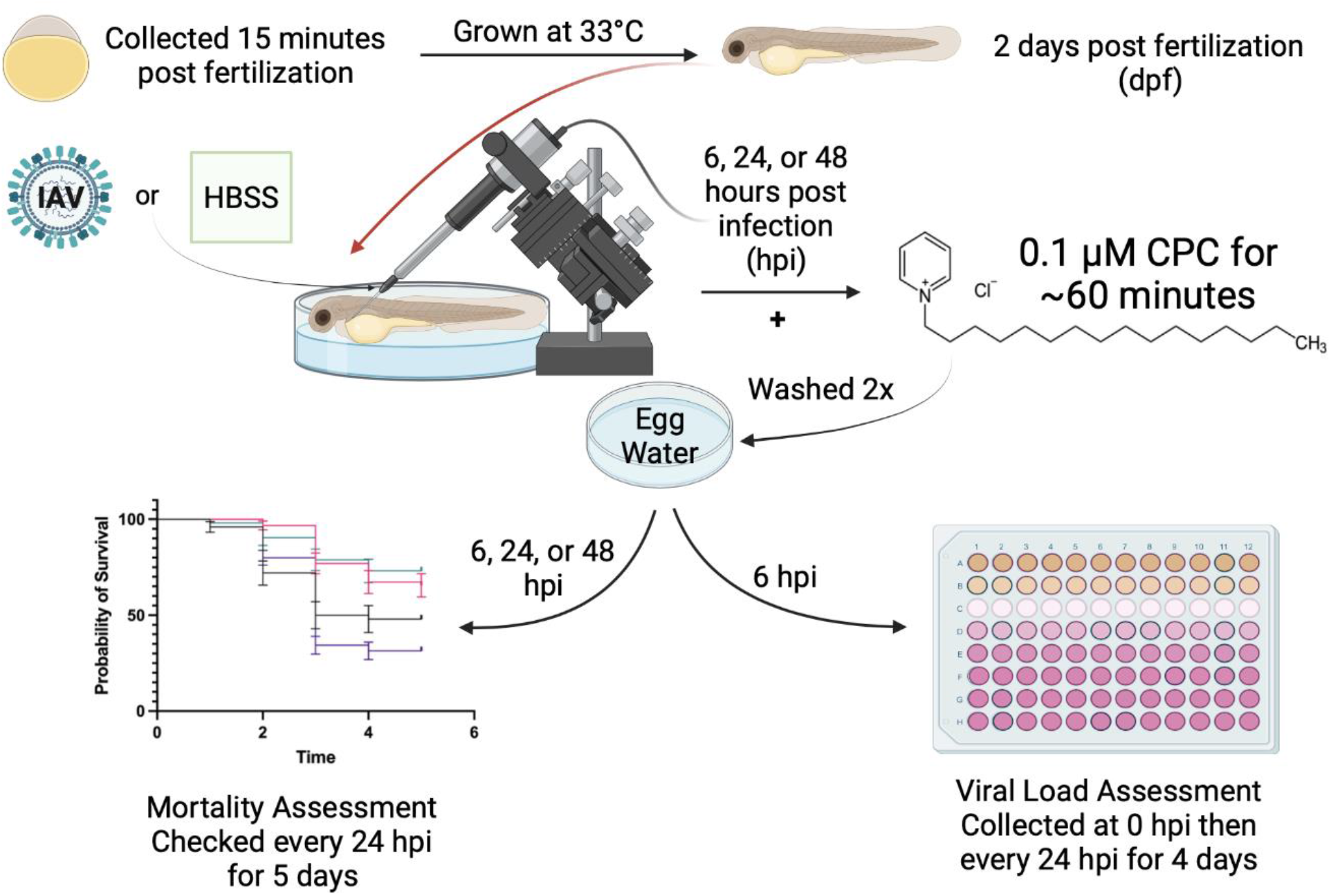
Zebrafish influenza infection and CPC exposure workflow. Zebrafish embryos were collected 15 minutes post-fertilization and grown at 33 °C. At 48 hours post fertilization (hpf), embryos were dechorionated (spontaneously or manually) and injected with IAV. CPC treatment of 0.1 µM in egg water was administered to the embryos at 6, 24, or 48 hours post infection (hpi) for a duration of ∼60 minutes before being washed two times to remove remaining traces of CPC from the egg water. Mortality and viral load were assessed every 24 hours for five days post-infection (dpi).

### Microinjection of IAV

Microinjections of influenza were performed as in (Gabor *et al.*, 2014), with the following modifications. Agarose gel (2%) was utilized in 100-mm petri dishes coated with 3% methylcellulose. After anesthetizing the embryos with sodium bicarbonate-buffered MS-222 (200 mg/L), a volume of 1.0 nl (∼ 8.7 ×10^2^ EID50) APR8 IAV or HBSS (Hank’s Balanced Salt Solution, Gibco) was microinjected. The control zebrafish were sibling zebrafish injected simultaneously with treated zebrafish. Needles containing IAV were changed every hour to guarantee the virus remained viable. Following injection, 45 zebrafish were sorted into plates of 50 mL egg water, which was changed daily. Pulled microcapillary pipettes (1.0 mm outside diameter, 0.6 mm inside diameter, Sutter Instruments, Novato, CA) were used to inject either the virus or HBSS (Goody *et al.*, 2014; Sullivan *et al.*, 2017).

### CPC Treatment on Zebrafish

CPC was administered to influenza infected and control, non-infected zebrafish at a concentration of 0.1 µM in 50 mL of egg water. CPC doses significantly higher than 0.1 µM caused embryo mortality (data not shown). The selection of the infected zebrafish to receive CPC treatment was random. The fish were treated with CPC for 1 hour at 33 °C under light-sensitive conditions (placed in a dark incubator). The fish were then rinsed twice in egg water to remove all traces of CPC and placed into fresh egg water for survivability experiments. Standard CPC treatment occurred at 6 hpi. Specialized treatments were administered 24, 48, and 72 hpi. Each timed CPC experiment was compared to CPC-treated zebrafish and to untreated zebrafish, for both IAV- and HBSS-injected zebrafish (Figure 10).

### TCID_50_ Assay

The burden of influenza virus in IAV-injected zebrafish was quantified using a TCID_50_ assay. Wild-type zebrafish embryos (150 per experimental group) were injected and maintained as described in “Microinjection of IAV.” Water was changed daily, and 25 fish from each group were collected, starting from 0 to 96 hpi. Embryos were placed into a kill tricaine solution (200 µg/mL) for 5 minutes to ensure death before being placed in a sterile microcentrifuge tube. The residual tricaine solution was removed and replaced with 500 μL of RNAlater (ThermoFisher Scientific) and flash frozen with liquid nitrogen. Samples were stored at −80 °C.

MDCK-London cells were plated in 96-well plates (USA Scientific) the night before the TCID_50_ assay, with eight wells for samples and with one well at the end of each row dedicated to the control, at a cell density designed to achieve 90% confluency on the day of the experiment. Each sample well was plated in triplicate to allow for three tests of the serial dilutions to confirm the viral concentration. In addition, cells were not allowed to grow past passage 10. Different passages and aliquots were utilized according to the cells’ difference in behavior, such that for experiment 1 there were 17,000 cells per well, experiment 2 had 13,000 cells per well, and experiment 3 was 14,500 cells per well. On the day of the experiment, cells were washed twice with 1x dPBS, pH 7.4, while the virus-containing samples were prepared as follows.

The RNAlater samples were thawed at room temperature, then RNAlater was removed manually with a Pasteur pipette and replaced with 1 mL of MEM-BSA/TPCK. A metal bead was added, and the samples were homogenized with a bullet blender at setting 3 for 5 minutes at 4 °C. The samples were briefly centrifuged at 8000 X g for 1 minute. The samples were placed on ice and moved to the BSL-2 hood. Serial dilutions of the samples were prepared, 1:10 until 10^-7^. The second dPBS wash was removed from the MDCK-London cells, and 50 µL of each sample dilution was added to corresponding plates in triplicate. Controls also received 50 µL of MEM-BSA/TPCK. Plates were centrifuged at room temperature for 5 minutes, 2000 X g before being placed in the 37 °C 5% CO_2_ incubator for 90 minutes. The samples were tapped every 15 minutes to ensure the cells did not dry out. Sample media was removed from the MDCK-London cells and 105 µL of MEM-BSA/TPCK was added to each well. Plates were grown at 37 °C with 5% CO_2_ for 3 days before being counted. Cells were visually assessed for cell damage such as death, morphological changes, and clearance; images were recorded and results were confirmed by a second researcher. The TCID_50_ was calculated by the Spearman-Kärber method (Hierholzer and Killington, 1996).

### Statistical Analyses

Statistical significance was determined by one-way ANOVA with a Tukey’s post-test for both the Trypan blue and LDH assays. Significance testing of FPALM images involved one-way ANOVA followed by Dunnett’s post-test performed using GraphPad Prism using data imported from the analyzed Matlab files. Standard curve comparison, a triplicate statistical analysis (Mantel-Cox, Log-rank, and Gehan-Breslow-Wilcoxon) in GraphPad Prism was used to determine significance in zebrafish survival studies. Two-way ANOVA with post Spearman’s correlation and normality tests of TCID_50_ assays in GraphPad Prism was performed to measure significance.

## Results

### CPC displaces PIP_2_-binding proteins MARCKS and PAmKate-PH from the plasma membrane

To test our hypothesis that CPC biochemically targets the negatively-charged eukaryotic lipid PIP_2_ and, therefore, interferes with PIP_2_-protein interactions, we assessed CPC effects on the cellular localization of the PIP_2_-binding protein MARCKS. RBL-2H3 mast cells were transiently transfected with monomeric red fluorescent protein (mRFP)-MARCKS-ED (Gadi *et al.*, 2011) and incubated overnight. On the next day, the cells were treated with Control buffer (0 µM CPC), 5 µM CPC, or 10 µM CPC for 30 minutes at 37°C, and live-cell images were collected with confocal microscopy (Figure 2A). The ratio of mean fluorescence intensity per pixel of (mRFP)-MARCKS-ED at the PM to the mean fluorescence intensity per pixel of (mRFP)-MARCKS-ED in the cytoplasm were calculated using Fiji ImageJ, and the ratio was calculated: “PM/cytosol mean fluorescence per pixel.” This ratio dose-responsively decreases with increasing CPC treatment (Figure 2B): 4.9 ± 0.2 for Control (0 µM CPC) samples, 2.4 ± 0.1 for 5 µM CPC samples, and 1.8 ± 0.1 for cells exposed to 10 µM CPC. Comparison of this ratio in CPC-treated and Control cells shows that CPC statistically significantly displaces the PIP_2_-binding protein MARCKS from the PM, into the cytosol (Figures 2A and 2B).

**Figure 2.**
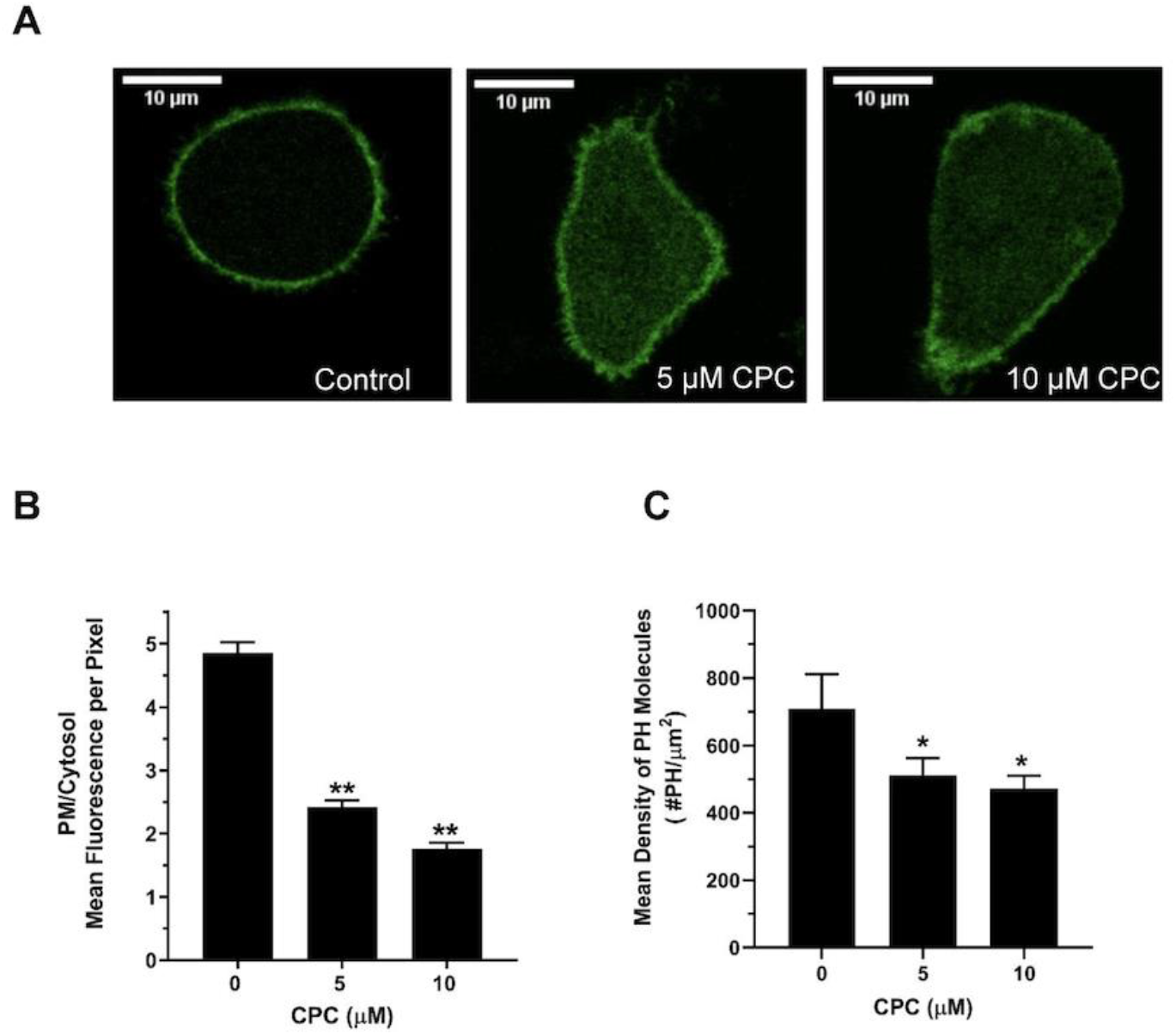
CPC reduces the PM/Cytosolic ratio of PIP_2_-binding protein MARCKS in RBL-2H3 cells and reduces the mean density of the PIP_2_-binding protein PAmKate-PH in PM of NIH-3T3 cells. (**A)** Representative live-cell confocal microscopy images of RBL-2H3 cells expressing mRFP-MARCKS-ED, ± CPC exposure: Control (0 µM CPC), 5 µM CPC, or 10 µM CPC for 30 minutes. After washing off the CPC with BT, confocal images were taken. (**B)** The ratio of the mean fluorescence per pixel of MARCKS at the PM and at the cytosol. Values represent mean ± SEM of three independent experiments that were derived from analysis of n = 143 cells for Control (0 µM CPC), n = 90 cells for 5 µM CPC, and n = 65 cells for 10 µM CPC. **(C)** Quantification of PH levels at the PM using super-resolution microscopy. NIH-3T3 cells expressing PAmKate-PH, ± CPC exposure for one hour, then fixed by 4% PFA. FPALM imaging of the PM of the transfected cells was carried out with TIRF excitation. The mean density of PH molecules was plotted as a function of CPC treatment. Values represent mean ± SEM of three independent experiments from analysis of n = 32 cells for BT Control (0 µM CPC), n = 30 cells for 5 µM CPC, and n = 31 cells for 10 µM CPC. Statistically significant results, as compared to Control (0 µM CPC), are represented by *p < 0.05 and **p < 0.01, as determined by one-way ANOVA followed by Dunnett’s post-test.

To test the same hypothesis, in an alternative system (different cell line, PIP_2_-binding protein, and microscopy method), we assessed CPC effects on the overall mean density at the plasma membrane of the PIP_2_-binding protein PAmKate-PH. NIH-3T3 cells were transfected with PAmKate-PH and another putative PIP_2_-binding protein HA-Dendra2 (Curthoys *et al.*, 2019). The next day, super-resolution imaging of the PM of the transfected cells was carried out with TIRF illumination to focus on the plasma membrane, excluding the cytoplasm. Cells were treated with Control buffer (0 µM CPC), 5 µM CPC, or 10 µM CPC for 1 hr at 37°C, cells were fixed, and images were collected with FPALM. The mean density of PAmKate-PH (“PH”) molecules was quantified by taking the ratio of the total number of PH molecules (PH localizations) over the cell area imaged in the same acquisition (following drift correction and bleedthrough correction), and plotted as a function of CPC treatment (Figure 2C). The mean density of PH molecules (# PH/μm^2^) was observed to be 700 ± 100 /µm^2^ for the Control (0 µM CPC) samples, 510 ± 50/µm^2^ for 5 µM CPC samples, and 470 ± 40/µm^2^ for cells exposed to 10 µM CPC. Comparison of the density of PH (# PH/μm^2^) in CPC-treated and Control cells shows that CPC significantly reduces the density of the PIP_2_-binding protein PAmKate-PH present adjacent to the PM (Figure 2C).

### CPC inhibits a PIP_2_-dependent cellular function of mast cells, degranulation

Degranulation in mast cells is the culmination of a PIP_2_-dependent signaling process (Santos *et al.*, 2013). Thus, to test our hypothesis that CPC biochemically targets PIP_2_ and thus interferes with PIP_2_-dependent cellular functioning, degranulation was measured following CPC exposure using the fluorescence assay described in (Weatherly *et al.*, 2013), adapted for the use of CPC. In this assay, cells were sensitized with anti-dinitrophenyl (DNP) mouse IgE, to occupy surface IgE receptors, which were then crosslinked with multivalent DNP-BSA antigen (Ag) to initiate cellular signaling. The dose of Ag utilized, 0.0004 μg/mL, was chosen because it elicited a moderate absolute degranulation response of 33% ± 1% (SEM) of the maximal possible granule release when in the absence of CPC. By choosing a moderate Ag dose, potential stimulatory or inhibitory CPC effects can be observed.

Figure 3A displays the effects of CPC on degranulation of IgE-sensitized RBL-2H3 mast cells incubated for 1 h in BT containing 0.0004 μg/mL Ag. CPC significantly inhibits degranulation in a dose-responsive manner, beginning with ∼25% inhibition at 1 µM (Figure 3A).

**Figure 3.**
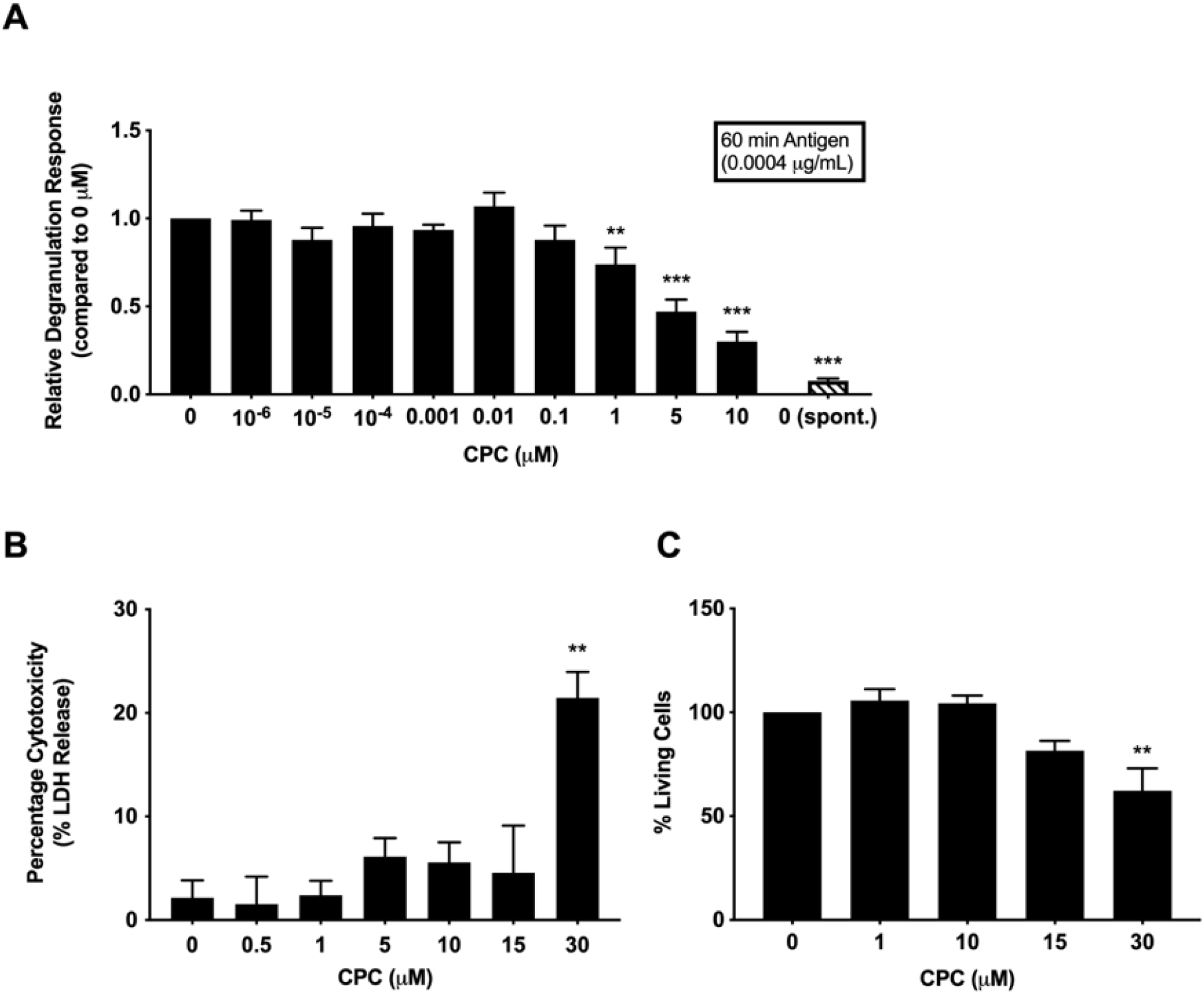
Non-cytotoxic doses of CPC inhibit RBL-2H3 mast cell degranulation. **(A)** RBL-2H3 mast cells were sensitized with IgE and stimulated to degranulate with 0.0004 μg/mL Ag for 1 hour, **±** CPC in BT; the spontaneous (“spont.”) response was determined by cells which were not exposed to Ag or CPC. Normalized degranulation response is plotted against CPC exposure concentrations. Effects of CPC on RBL-2H3 cell survivability were assessed by LDH assay **(B)** and by trypan blue-exclusion **(C)**. Values presented are means ± SEM of three independent experiments; three replicates were used in each experiment. Statistical significance, compared to the Control (0 µM CPC), is represented by **p < 0.01, ***p < 0.001 and determined by one-way ANOVA followed by Tukey’s post-test.

Statistically significant inhibition of degranulation began at 1 μM CPC after 1 hour exposure, resulting in a degranulation response that was 0.7-fold ± 0.1 (SEM) of the 0 μM CPC Control level (Figure 3A). At 5 μM CPC, degranulation was reduced to 0.47-fold ± 0.07 (SEM) of the 0 μM CPC Control response. At 10 μM CPC, the response was reduced to 0.30-fold ± 0.05 (SEM) of the 0 μM CPC Control, equivalent to a 70% inhibition of this cellular function. We confirmed that this inhibition was a true cellular effect, not due to interference of CPC with the reaction of β-hexosaminidase with its fluorogenic substrate, which is the signal used to quantify degranulation; also, CPC does not directly affect fluorescence of the background signal (Figure S1). These data show that CPC inhibits mast cell degranulation in a dose-responsive fashion.

### CPC is not cytotoxic to RBL-2H3 cells at concentrations that inhibit degranulation and displace MARCKS

Two cytotoxicity assays were conducted to assess the survivability of RBL-2H3 cells when exposed to a range of CPC doses for 1 hour. To assess the survivability of RBL-2H3 cells upon exposure to various CPC doses, we performed a lactate dehydrogenase (LDH) release assay. LDH is an intracellular enzyme released upon plasma membrane damage or death. LDH release was not significantly altered, compared to 0 μM CPC Control, for exposures up to 15 μM CPC for 1 hour, indicating a lack of cytotoxicity or plasma membrane damage (Figure 3B); however, 30 μM CPC for 1 hour caused LDH release. Activity of purified LDH was unaffected by CPC, indicating that CPC does not interfere with the detection mechanism of the LDH assay itself (Figure S2).

A trypan blue-exclusion assay was used to assay metabolic viability of RBL-2H3 mast cells upon 1 hour CPC exposure. Figure 3C shows a statistically significant cytotoxic effect only in cells exposed to 30 μM CPC, while concentrations ≤ 15 μM were not cytotoxic as quantified by the assay. Furthermore, in NIH-3T3 cells, there was a lack of CPC cytotoxicity at the doses tested (5 and 10 μM) in a trypan blue assay (Figure S3).

Taken together, the results from these two assays strongly suggest that CPC is not cytotoxic to RBL-2H3 and NIH-3T3 cells when exposed for 1 hour at concentrations ≤ 15 μM. These data suggest that cell death is not the mechanism for the effects of CPC documented in this manuscript (such as degranulation and re-distribution of PIP_2_-binding proteins).

### CPC disrupts clusters of the influenza viral protein hemagglutinin in NIH-3T3 cells

Considering the relationship between PIP_2_ and the influenza viral protein hemagglutinin (HA) (Curthoys *et al.*, 2019), the requirement of HA clustering for influenza infectivity (Ellens *et al.*, 1990; Takeda *et al.*, 2003), the reported antiviral properties of CPC (Popkin *et al.*, 2017), and the disruption of PIP_2_-binding protein MARCKS by CPC, we employed super resolution microscopy to test whether CPC affects nanoscale HA clustering. We used FPALM (Hess *et al.*, 2006) with TIRF excitation to image the PM of NIH-3T3 cells expressing both HA-Dendra2 and PAmKate–PH Domain, which binds to and labels PIP_2_ (Gambhir *et al.*, 2004; Curthoys *et al.*, 2019). Clusters were identified using SLCA (see methods). Quantification of HA clusters showed a wide range of density with the mean value of 12700 ± 4300 HA/µm^2^ for control (Figure 4A). This wide variation of density in HA cluster was absent in cells treated with CPC: the mean HA density in CPC-treated cells decreased significantly by **_∼_**79% when compared against the control, with the mean HA density of 2700 ± 400 HA/µm^2^ for 5 µM and 2700 ± 400 HA/µm^2^ for 10 µM CPC-treated cells (Figure 4A). Similarly, the number of HA molecules in a cluster was reduced significantly by CPC. On average, 340 ± 110 HA molecules were found in an HA cluster which decreased significantly by **_∼_**74% and **_∼_**76% to 88 ± 9 HA molecules and 81 ± 13 HA molecules for 5 µM and 10 µM CPC-treated cells, respectively (Figure 4B). The mean area of an HA cluster was observed to be 0.041 ± 0.004 µm^2^ for Control, 0.044 ± 0.006 µm^2^ for 5 µM, and 0.039 ± 0.003 µm^2^ for 10 µM CPC (Figure 4C). The mean perimeter of an HA cluster was observed to be 0.90 ± 0.08 µm for control 1.0 ± 0.1 µm for 5 µM, and 0.09 ± 0.06 µm for 10 µM CPC (Figure 4D). Both the mean area and mean perimeter of an HA cluster were statistically unaffected by CPC treatment. The mean density of HA molecules (within each cell) was quantified by dividing the total number of HA molecules (HA localizations) in a given cell over the imaged area of each cell. The mean density was 195 ± 8 HA /µm^2^ in the Control, 190 ± 12 HA/µm^2^ for 5 µM, and 180 ± 12 HA/µm^2^ for 10 µM CPC treatment: no statistically significant CPC effect was observed.

**Figure 4.**
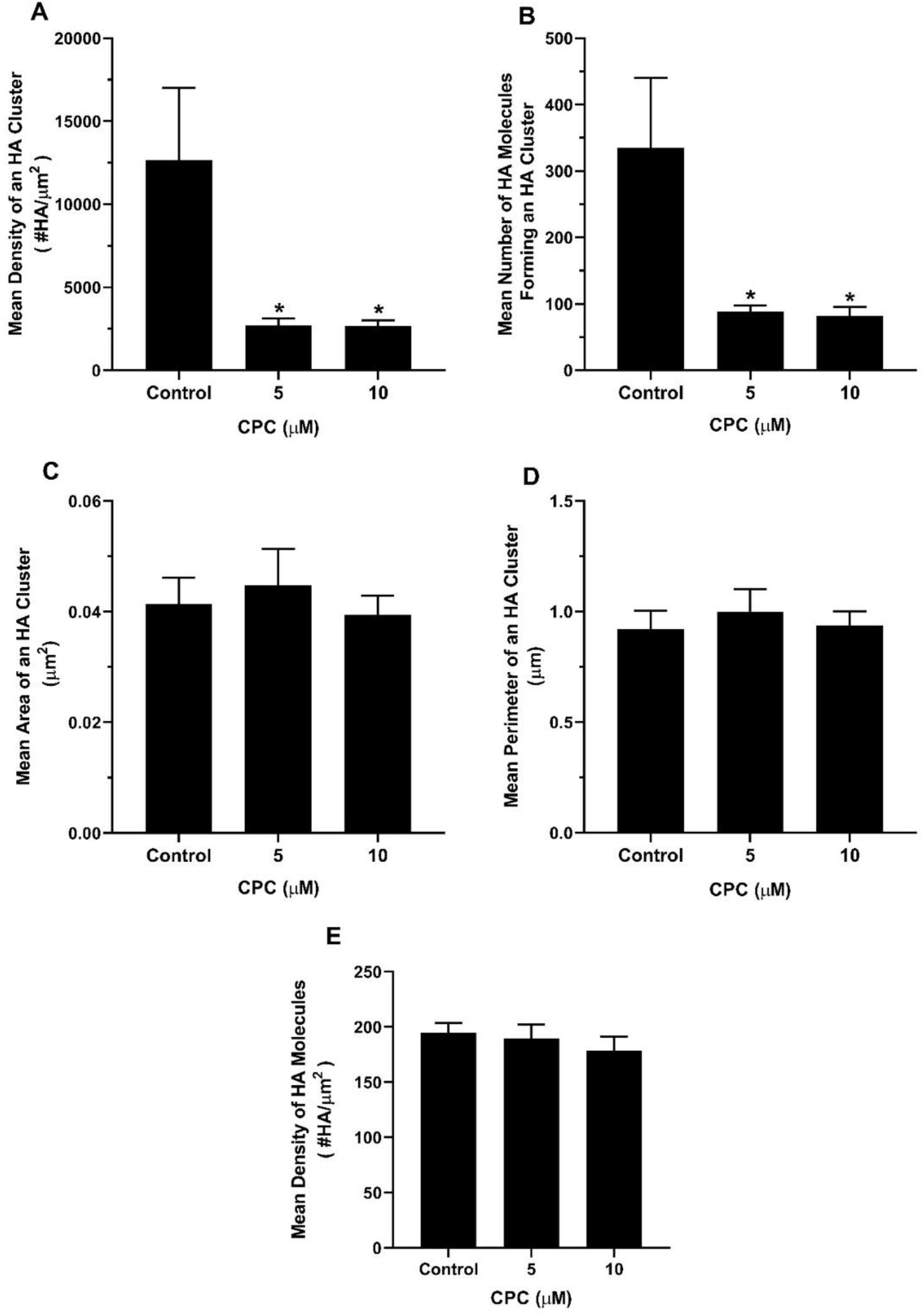
CPC alters HA cluster properties in NIH-3T3 cells. Two-color TIRF FPALM was used to obtain images from fixed NIH-3T3 cells co-expressing HA-Dendra2 and PAmKate-PH constructs. NIH-3T3 cells were exposed to Control (0 µM CPC), 5 µM, and 10 µM CPC for 1 hour at 37 °C, then were fixed with 4% PFA. HA cluster properties **(A)** Mean density of an HA cluster **(B)** Mean number of HA molecules forming an HA cluster **(C)** Mean Area of an HA cluster **(D)** and Mean Perimeter of an HA cluster were quantified using SLCA. **(E)** The mean density of HA molecules in each cell was quantified by taking the ratio of the total number of HA molecules (HA localizations) over the cell area. Values represent mean ± SEM of three independent experiments from analysis of n = 24 cells for BT Control (0 µM CPC), n = 20 cells for 5 µM CPC, and n = 22 cells for 10 µM CPC. Statistically significant results are represented by *p < 0.05, as compared to Control by one-way ANOVA followed by Dunnett’s multiple comparison test against the Control.

### CPC disrupts PIP_2_ clusters in NIH-3T3 cells

We also studied the effect of CPC in PIP_2_ clusters along with the HA, as PIP_2_ has also been previously reported to cluster at the PM (Curthoys *et al.*, 2019) (van den Bogaart *et al.*, 2011; Wang and Richards, 2012). In order to visualize the nanoscale distribution of PIP_2_, we used FPALM to image a PAmKate-tagged version of the PIP_2_-binding Pleckstrin Homology (PH-) domain from PLC-δ (Gambhir *et al.*, 2004; Curthoys *et al.*, 2019), called PAmKate-PH and abbreviated as “PH domain” (Figure 5). Cluster analysis was performed using SLCA (see Methods). The mean density of a PH domain cluster was 2700 ± 500 /µm^2^ for Control, 2400 ± 300 /µm^2^ for 5 µM, and 2100 ± 100 /µm^2^ for 10 µM CPC treated cells (Figure 5A); no statistically significant CPC effect was detected on mean density. However, the mean number of PAmKate-PH molecules forming a cluster decreased significantly (p < 0.01). On average, 270 ± 30 molecules formed a PH domain cluster (Control) while only 170 ± 13 and 190 ± 21 PH domain molecules on average were found in a cluster for 5 µM and 10 µM CPC-treated cells, respectively (Figure 5B). The mean area of a PH domain cluster (Control) was found to be 0.105 ± 0.009 µm^2^, which appeared to decrease to 0.08 ± 0.01 µm^2^ and 0.085 ± 0.006 µm^2^ for 5 µM CPC and 10 µM CPC-treated cells, respectively (Figure 5C); however, no statistical significance was observed by one-way ANOVA followed by Dunnett’s multiple comparison test against Control, and the Kruskal-Wallis test was barely insignificant (p = 0.0503). The mean perimeter of PH domain clusters was statistically unchanged by CPC: 1.7 ± 0.1 µm for Control, 1.4 ± 0.1 µm for 5 µM, and 1.53 ± 0.08 µm for 10 µM CPC (Figure 5D).

**Figure 5.**
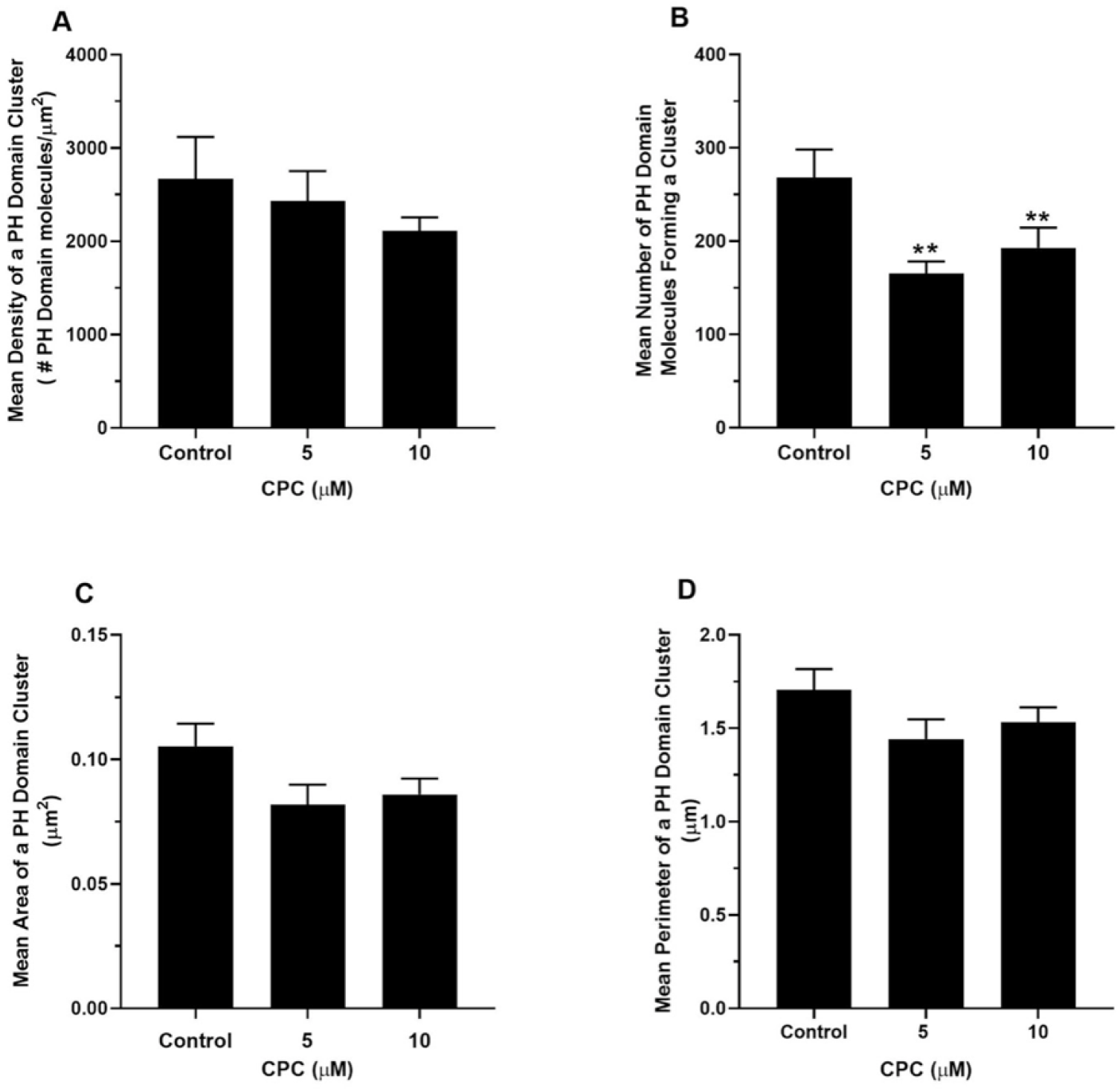
CPC affects PIP_2_-binding protein cluster properties in NIH-3T3 cells. Two-color TIRF FPALM was used to obtain images from fixed NIH-3T3 cells co-expressing HA-Dendra2 and PAmKate-PH. NIH-3T3 cells were exposed to Control (0 µM CPC), 5 µM, and 10 µM CPC for 1 hour at 37 °C, then were fixed with 4% PFA. PH domain cluster properties **(A)** Mean density of a PH domain cluster, **(B)** Mean number of PH domain molecules forming a cluster, **(C)** Mean Area of a PH domain cluster, **(D)** Mean perimeter of a PH domain cluster for Control, 5 µM, and 10 µM CPC-treated cells were quantified using SLCA. Values represent mean ± SEM of three independent experiments from analysis of n = 25 cells for BT Control (0 µM CPC), n = 26 cells for 5 µM CPC, and n = 29 cells for 10 µM CPC. Statistically significant results are represented by **p < 0.01, as compared to the Control by one-way ANOVA followed by Dunnett’s multiple comparison test against the Control.

### CPC reduces the co-clustering of hemagglutinin and PH Domain in NIH-3T3 cells

CPC dramatically reduced colocalization of HA-Dendra2 and PAmKate-PH in NIH-3T3 cells (Figure 6). In order to quantify the effect of CPC on HA and PH domain co-clustering at the nanoscale, the mean number of HA molecules per grid pixel within PH clusters (N_HA-PH_; see Methods) and the mean number of PH molecules within HA clusters (N_PH-HA_) were determined from two-color FPALM results. The analysis shows a significant decrease in N_PH-HA_ in 10 µM CPC-treated cells, decreasing by approximately 71% as compared to control, which was statistically significant (Figure 7A). A similar result was obtained for the number of HA localizations within PH domain clusters, where the Mean Pixel Sum N_HA-PH_ dropped approximately by 61% compared to Control, again statistically significant (Figure 7B).

**Figure 6.**
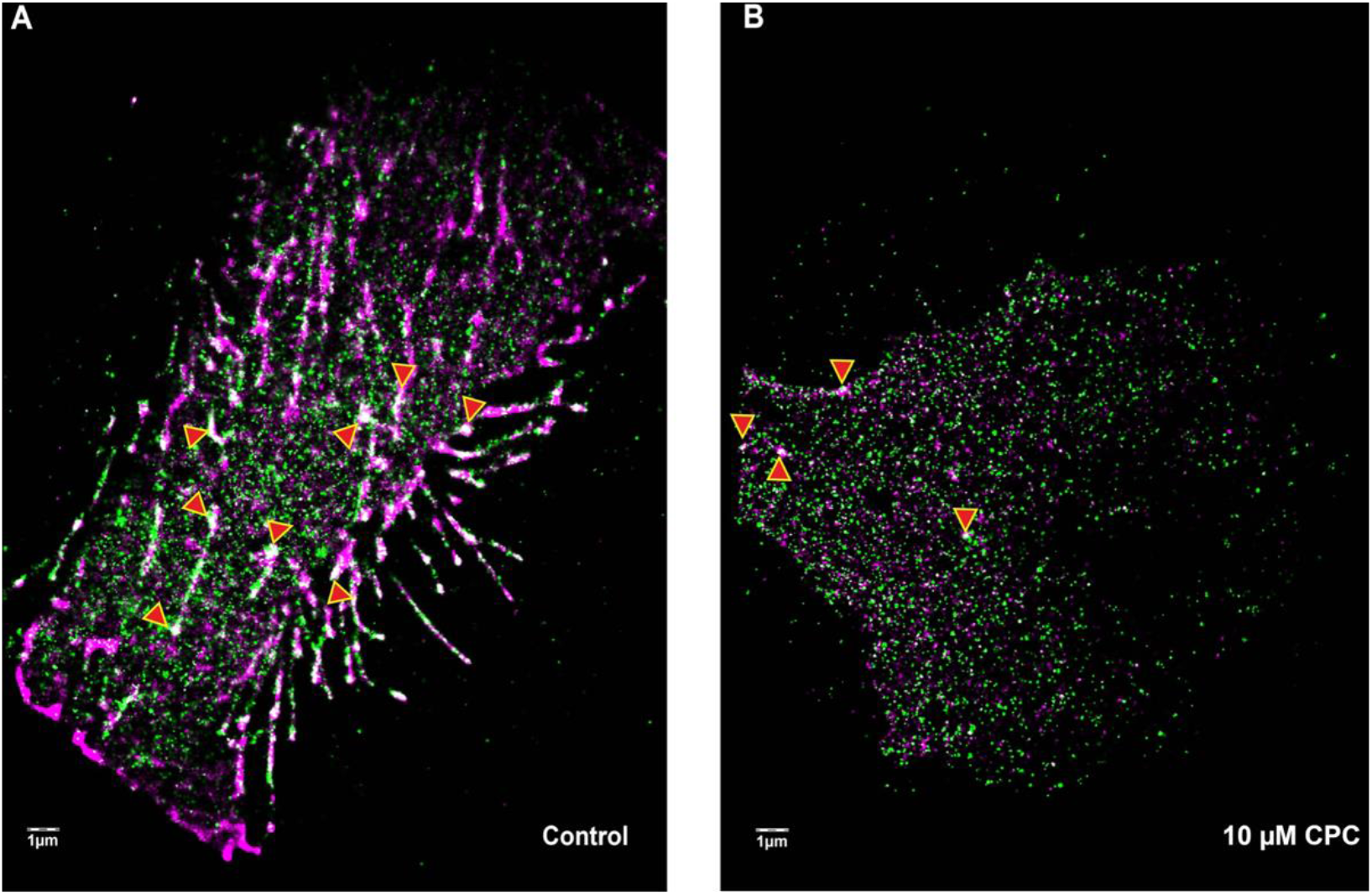
CPC disrupts nanoscale colocalization of HA-Dendra2 and PAmKate-PH imaged in fixed NIH-3T3 cells. Two-color FPALM images of fixed NIH-3T3 cells expressing HA-Dendra2 (green) and PAmKate-PH (magenta) of **(A)** Control and **(B)** 10 µM CPC-treated cells. Arrows (red with yellow outline) point to areas of colocalization (white).

**Figure 7.**
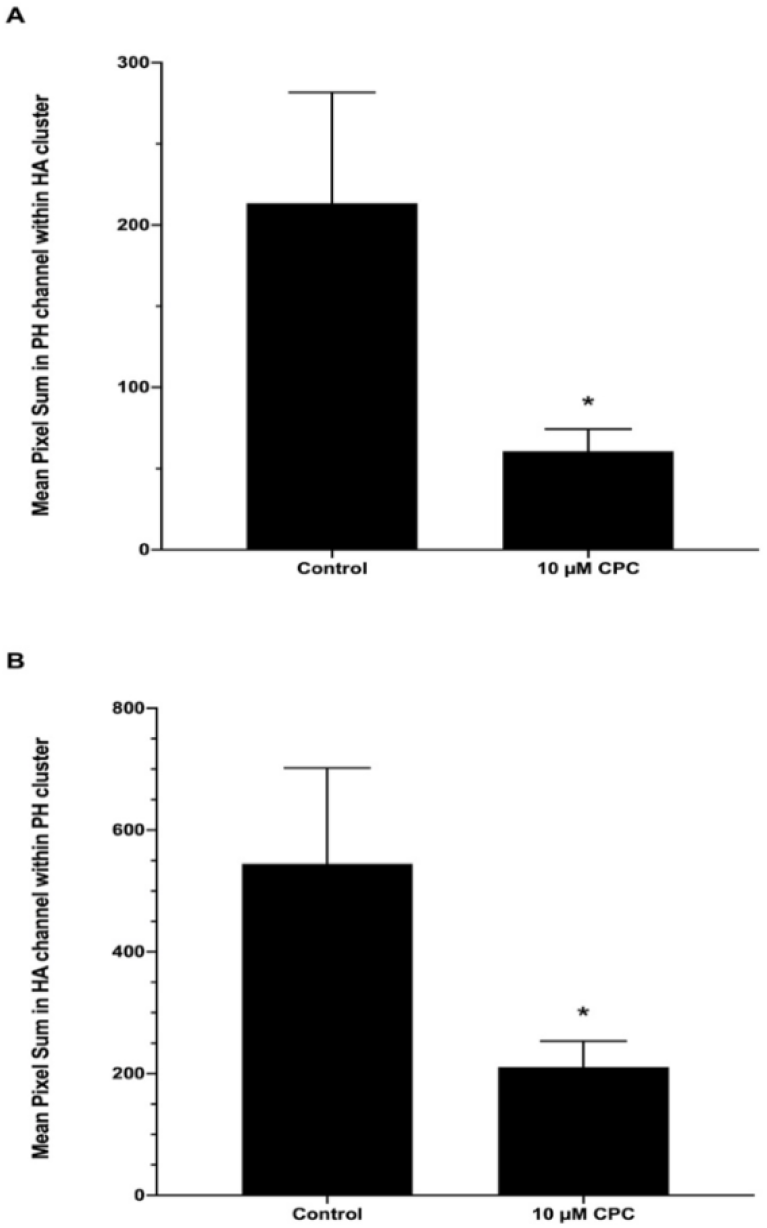
CPC alters co-clustering of HA-Dendra2 and PAmKate-PH in NIH-3T3 cells. Co-clustering was quantified from the distribution of Dendra2-HA and PAmKate-PH molecules localized within a grid of 35 nm x 35 nm pixels, after bleed-through correction, in fixed NIH-3T3 cells. **(A)** Pixels identified as HA-Dendra2 clusters (containing at least 5 HA localizations) were selected. The number of PAmKate-PH localizations was then summed over all pixels within the selection, and then averaged over all cells, and is plotted as “mean pixel sum” (see Methods) as a function of CPC treatment (10 µM). **(B)** Pixels identified as PAmKate-PH clusters (at least 5 PH domain localizations) were selected. The number of HA-Dendra2 localizations was then summed over all pixels within that selection and then averaged over all cells and is plotted as “mean pixel sum” as a function of CPC treatment (10 µM). Values represent mean ± SEM of three independent experiments from analysis of n = 31 cells for BT Control (0 µM CPC) and n = 30 cells for 10 µM CPC. Statistically significant differences are represented by *p < 0.05 as compared to Control, determined by unpaired t-test.

### CPC treatment reduces influenza infections and increases survival in AB zebrafish embryos

Viral load and mortality assessment were performed ± CPC exposure at 0.1 μM for 1 hour in *Danio rerio* (zebrafish, ZF), an *in vivo* influenza virus infection (IAV) model. In order to determine appropriate dose and timing of CPC exposure, first, acute (1 hour) exposures of 2-4 dpf ZF embryos to CPC alone (no IAV injection) were performed (Figure 8). All doses 1 μM and above were 100% lethal to 2-4 dpf ZF embryos within 3 days post CPC exposure, while 0.5 μM and 0.25 μM CPC were lethal to a fraction of animals. In contrast, 99.5% of 200 exposed ZF embryos survived for at least 3 days following a 1 hour, 0.1 μM CPC exposure. Next, longer exposures to this chosen 0.1 μM CPC dose were performed, but lethality (15% of embryos) began within 3 hours of continuous CPC exposure, and mortality increased with length of CPC exposure (43% lethal at 6 hours of 0.1 μM CPC, 96% lethal at 12 hours, 100% lethal at 24 hours) (these data are not plotted). Thus, exposure to CPC at 0.1 μM for 1 hour exposure was chosen for anti-influenza pharmacological studies.

**Figure 8.**
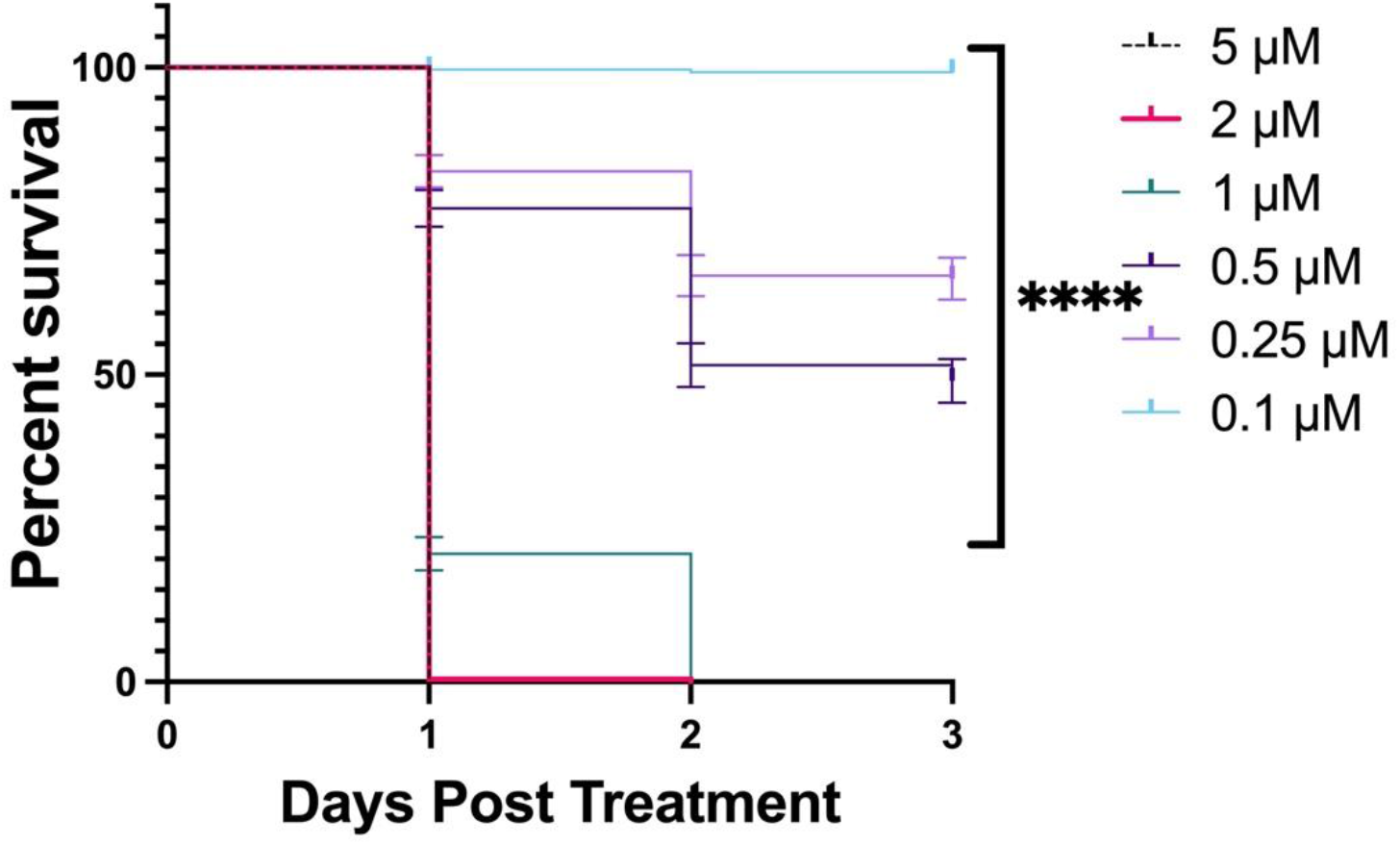
Cytotoxic doses of CPC for the extensive evaluation of zebrafish embryo survival. Zebrafish embryos injected with HBSS and exposed to various doses of CPC were monitored for cumulative survival over the course of 5 days (no observable death occurred after 3 days). CPC doses higher than 0.5 µM showed significant decrease in survival. Values presented are for a total of 204 zebrafish, assayed in four independent experiments. Statistically significant differences are represented by ****p < 0.0001, determined by the Mantel-Cox test, the Logrank test, and the Gehan-Breslow-Wilcoxon test of survival curves in GraphPad Prism.

Specifically, 1 hour of 0.1 μM CPC treatment in mild to moderate IAV infections was shown to significantly increase the chance of survival if administered within the first 48 hpi (Figure 9). For zebrafish exposed for 1 hour to 0.1 μM CPC at 6 hpi, survival was analyzed at 5 dpi: average survival of CPC-treated zebrafish was 83.2%, versus 60.1% for CPC-untreated, IAV-infected zebrafish (Figure 9A). For zebrafish exposed for 1 hour to 0.1 μM CPC at 24 hpi, survival was analyzed at 5 dpi: average survival of CPC-treated zebrafish was 87.1%, versus 60.1% for CPC-untreated, IAV-infected zebrafish (Figure 9B). For zebrafish exposed for 1 hour to 0.1 μM CPC at 48 hpi, survival was analyzed at 5 dpi: the average survival of CPC-treated zebrafish was 87.7%, versus 60.1% for CPC-untreated, IAV-infected zebrafish (Figure 9C). Importantly, neither CPC nor mock HBSS injection causes mortality within 5 dpi (Figure S4). If the one-hour CPC treatment is withheld until 72 hpi, CPC is unable to rescue the embryos from influenza mortality (Figure S5).

**Figure 9.**
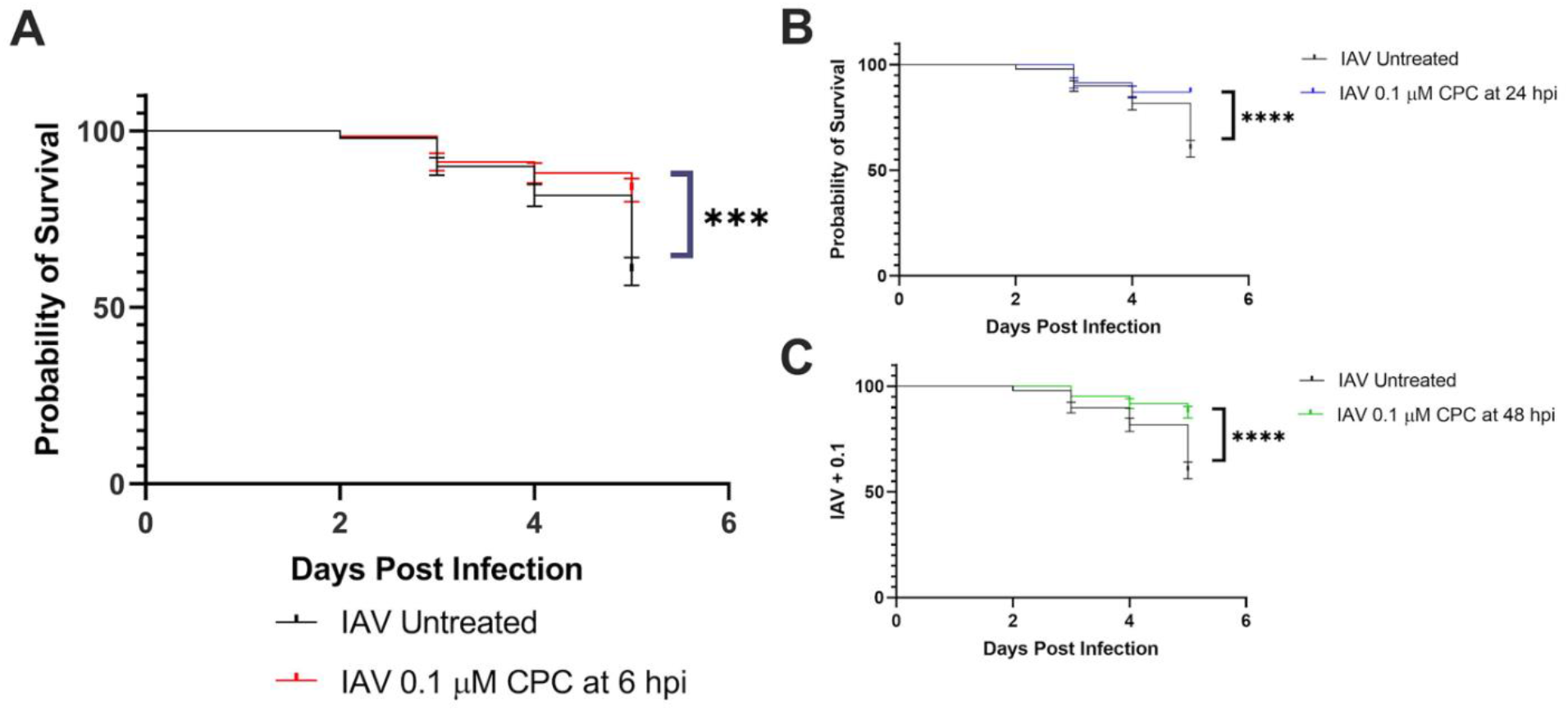
CPC increases zebrafish embryo survival rate following influenza infection. Zebrafish embryos (28 hpf) were injected with 1.0 nl (∼ 8.7 ×10^2^ EID50) of APR8 IAV into the duct of Cuvier, for all groups in this figure. Survival curves of IAV-infected zebrafish, ± 0.1 µM CPC treatment for one hour at 6 hpi **(A)**, 24 hpi **(B), or** 48 hpi **(C).** Values presented are for a total of ∼150 zebrafish, assayed in three independent experiments. Statistically significant differences are represented by ***p < 0.001, ****p < 0.0001, determined by the Mantel-Cox test, the Logrank test, and the Gehan-Breslow-Wilcoxon test of survival curves in GraphPad Prism.

This CPC reduction in influenza mortality correlates to CPC reduction in viral burden despite a similar starting viral load (Figure 10). Figure 10 shows that there is no difference in viral burden (TCID_50_/mL) at 0 hpi for IAV-infected zebrafish with or without the CPC treatment (1 hour, 0.1 μM). However, starting at 24 hpi, and especially significant at 48 hpi, there is a decrease in viral burden by as much as 1 log-fold due to CPC treatment (Figure 10). Although it appears that viral burden drops, even in non-CPC treated fish, at the later time points (Figure 10), the reason is due to a combination of zebrafish mortality (fish that die due to influenza at earlier time points are not assessed at 96 hours), viral clearance, and removal of fish from the experiment.

**Figure 10.**
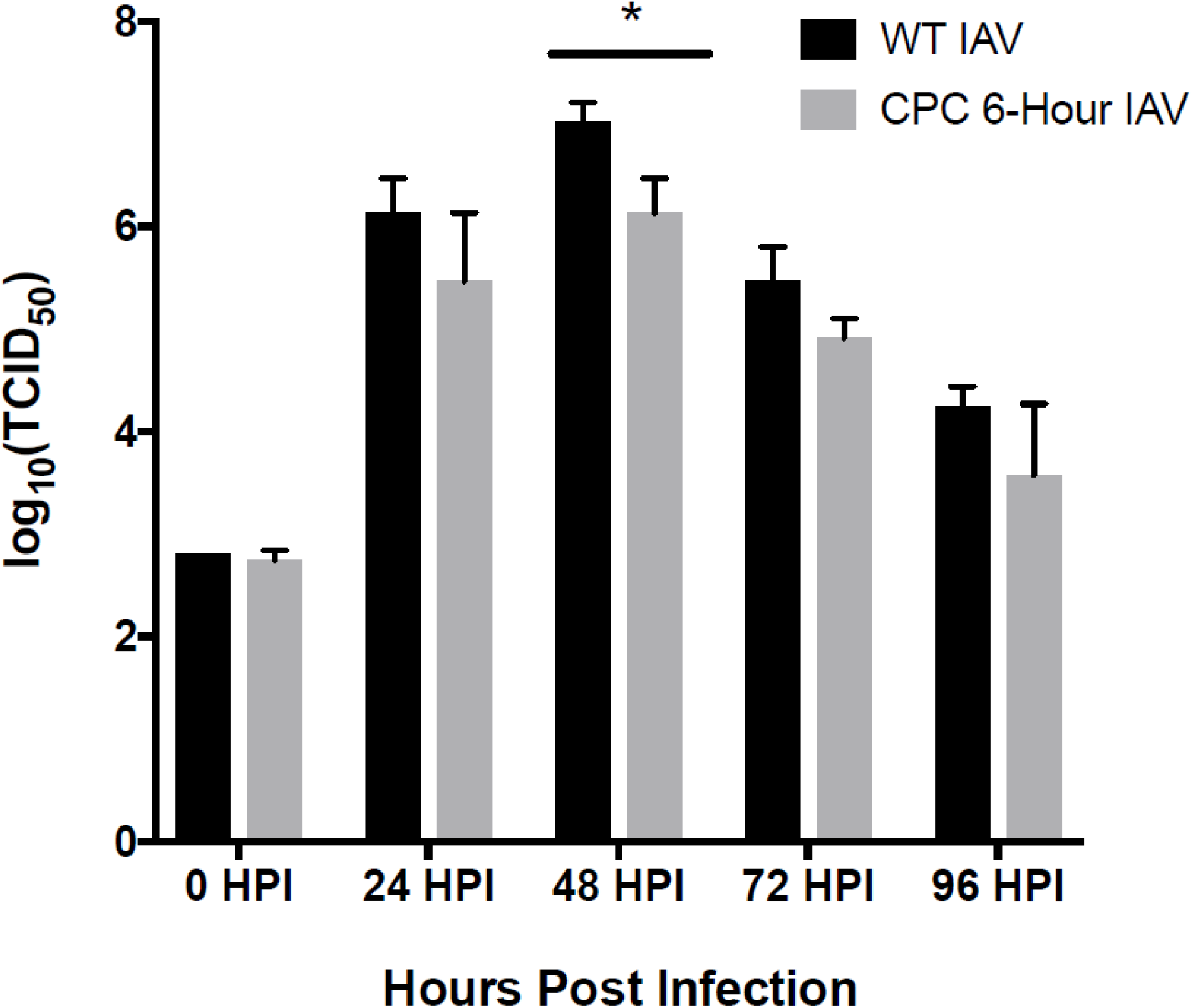
CPC treatment reduces viral load in AB zebrafish embryos. Zebrafish embryos (2 dpf) were injected with IAV as in Figure 9 (duct of Cuvier, 1.0 nl, ∼ 8.7 ×10^2^ EID50 of APR8 IAV). TCID_50_ comparison between groups with (green) and without (blue) 0.1 µM CPC administered for one hour at 6 hpi, in three replicate experiments. Statistical analysis of the difference between CPC-treated and untreated yielded p < 0.0005 using a two-way ANOVA.

## Discussion

Cetylpyridinium chloride possesses antiviral activity, including against influenza, in human (Mukherjee *et al.*, 2017), mouse (Popkin *et al.*, 2017), and, now, zebrafish with a short (1 hour), low-dose (0.1 μM) exposure (Figure 9). However, its anti-influenza mode of action, particularly at low-micromolar doses relevant to safe potential pharmacological usage, is unknown. At the CPC doses (≤ 10 μM) tested in zebrafish and cells in this study, CPC is far below its CMC (Mandal and Nair, 2002; Varade *et al.*, 2005; Abezgauz *et al.*, 2010; Shi *et al.*, 2011) and, thus, is not acting as a detergent (confirmed by lack of cytotoxicity, Figure 3B-C and lack of zebrafish mortality, Figure S4). Also, these low-micromolar doses are well below levels of CPC known to act as direct virucidal agents (Popkin *et al.*, 2017; Baker *et al.*, 2020; Koch-Heier *et al.*, 2021). Thus, because CPC is not acting via a detergent lysis or direct virucidal mechanism to alleviate zebrafish viral load and mortality following influenza infection, another anti-infective mechanism must be involved. In this study, the role for low-micromolar CPC effects on mammalian host cells and on interactions between the host cell and viral components was explored. We hypothesized that CPC affects the mammalian cell itself, to reduce influenza morbidity. We found that CPC disrupts interactions between PIP_2_ and multiple PIP_2_-binding proteins, including MARCKS, PH-domain from PLCδ, and influenza HA, suggesting a PIP_2_-focused mechanism underlying CPC disruption of influenza infectivity.

One of the central molecular players in mast cell signaling is PIP_2_. Thus, we hypothesized that, via its positive charge at the end of a lipidic tail, CPC biochemically targets the negatively-charged eukaryotic lipid PIP_2_ and, therefore, interferes with PIP_2_-protein interactions that are essential to mast cell function. These proteins could be MARCKS, PLC**γ**, or other PIP_2_-binding proteins that play crucial roles in mast cell function. Thus, we performed experiments to test this CPC mode-of-action hypothesis and found that CPC treatment caused MARCKS (in mast cells) and PH molecules (in NIH-3T3 cells), two well-known PIP_2_ binding proteins (McLaughlin *et al.*, 2002), to fall off the PM (Figure 2).

Concurrently, we discovered that CPC inhibits degranulation of antigen-stimulated mast cells in a dose-responsive manner, beginning at 1 µM (Figure 3A). These data represent one of the first studies of CPC effects on eukaryotic cells, despite its widespread use in personal care products. These findings indicate that further research into the effects of CPC on immune cell function are necessary and relevant to consumer health. The doses tested are non-cytotoxic, as indicated by the LDH and trypan blue cytotoxicity assays (Figure 3B-C); thus, the CPC effect on mast cells is not due to cell death. Thus, CPC is likely interfering with one or more components of the antigen-stimulated mast cell signaling cascade, resulting in its suppressive effect on degranulation. Further research will be necessary to pinpoint the mechanism of CPC disruption of mast cell degranulation.

Our results suggest that other PIP_2_-dependent physiological processes, including influenza infection, may be altered by CPC. Because of a recently-identified relationship between HA and PIP_2_ in which PIP_2_ modulates HA clustering, HA modulates PIP_2_ clustering, and both molecules are co-localized at the nanoscale in the PM (Curthoys *et al.*, 2019), we hypothesized that CPC may also affect the plasma membrane distribution of HA, via PIP_2_ interference. Based on our previously published results (Curthoys *et al.*, 2019), we hypothesized that HA interacts with PIP_2_ through charge-charge interactions between the negatively-charged PIP_2_ head and the HA cytoplasmic tail (CT), which has two basic residues and an estimated net charge of +6 per trimer at physiological pH, and also has several conserved, acylated cysteines within the HA CT which could interact with lipids within the inner leaflet of the PM (Veit *et al.*, 2013). This information led us to hypothesize that CPC blocking of this HA-PIP_2_ interaction would affect HA clustering, which is crucial for infectivity (Ellens *et al.*, 1990) and the viral lifecycle (Takeda *et al.*, 2003). We used super-resolution microscopy (FPALM) to image HA-Dendra2 and the PIP_2_-binding PH domain tagged with PAmKate (Flesch *et al.*, 2005; Szentpetery *et al.*, 2009; Curthoys *et al.*, 2019) (Figure 6). Localized molecular coordinates of HA-Dendra2 and PAmKate-PH were then quantified using SLCA (Curthoys *et al.*, 2019; Sangroula *et al.*, 2020; Sneath, 1957) to determine the properties of clusters formed by the HA and PH domain.

The density of HA in clusters in control cells varied widely (Figure 4A), consistent with published results showing that HA clusters exist on many length scales (Hess *et al.*, 2005). Significant disruption of HA clusters by CPC was observed, particularly with respect to the density and the number of HA molecules forming each HA cluster in CPC-treated cells (Figure 4A-B). Since HA is a transmembrane protein, CPC is not expected to cause HA to dissociate from the PM as we observe with PH and MARCKS proteins (Figure 2), but rather is disrupting the clusters through redistribution of the HA which is present in the PM. We speculate that this disruption could be due to CPC interactions with PIP_2_ or through interference with the interaction between the HA CTD and the PIP_2_ headgroup. These data are important because dense HA clusters are crucial for efficient viral entry through membrane fusion (Takeda *et al.*, 2003), and CPC significantly reduces HA cluster density, thus illuminating a potential mechanism of antiviral properties of CPC at low concentrations.

This mechanism is also consistent with our findings that CPC treatment increases survival (Figure 9) and reduces viral load (Figure 10) in influenza-infected zebrafish. There is a minimal difference in effect of CPC treatments at 5 and 10 μM on HA clustering (Figure 4), suggesting that 5 μM CPC may be a sufficient cellular dose for maximizing anti-influenza CPC activity. At 72 hpi, the infection is too severe for CPC treatment to make a difference (Figure S5). CPC does seem to rescue older fish better than younger fish (data not shown), indicating possible CPC disruption of developmental processes. The zebrafish embryos were injected with 1.0 nl (∼ 8.7 ×10^2^ EID50) of APR8 IAV and selected at random for CPC treatment using 0.1 µM for 1 hour (standard CPC treatment at 6 hpi and specialized treatments at 24, 48, and 72 hpi). In contrast, in the Popkin study, wildtype mice were infected intranasally (LD50, 8.0 x 10^3^ pfu/mouse) with the mouse-adapted influenza strain A/PuertoRico/8/1934 H1N1 (PR8, ATCC) while CPC was given orally (5 μL, in formulation) 15 minutes prior and then twice a day for 5 consecutive days (Popkin *et al.*, 2017). While the influenza and CPC exposures were different for this zebrafish study and the published mouse study, both found that CPC improves survivability following influenza infection.

PIP_2_ is known to cluster at the PM (van den Bogaart *et al.*, 2011; Wang and Richards, 2012; Curthoys *et al.*, 2019), and the interactions of PIP_2_ with proteins control a large number of cellular functions (McLaughlin *et al.*, 2002; McLaughlin and Murray, 2005; Di Paolo and De Camilli, 2006; Balla, 2013; O’Donnell, 2018). Quantification of PAmKate-PH (PIP_2_) clusters visualized by FPALM showed significant disruption by CPC (Figure 5), thus further confirming the effect of CPC on PIP_2_-binding proteins. Results also uncovered a major disruption of HA-Dendra2 co-localization with PAmKate-PH (Figures 6 and 7), further confirming that CPC can disrupt interactions between PIP_2_ and proteins at the PM. These results are important because PIP_2_-binding proteins have been previously shown to have polybasic domain and clusters to negatively-charged PIP_2_ and PIP_3_ through charge-charge interaction (Heo *et al.*, 2006), and CPC disrupts the clustering of several of these proteins. The CPC-induced reduction of the mean number of PH domain molecules forming a cluster also explains the observation that the only PH cluster property that is affected is the mean number of PH molecules per cluster, which would be due to CPC ejecting or screening some of the PH-domain-containing proteins close to the plasma membrane.

In addition to interactions with HA, PIP_2_ plays numerous roles in modulating cytoskeletal function and membrane biochemistry that may be involved in influenza pathogenesis (Raucher *et al.*, 2000; Di Paolo and De Camilli, 2006; Guerriero *et al.*, 2006; Heo *et al.*, 2006; Altan-Bonnet and Balla, 2012). PIP_2_ regulates cytoskeleton-PM adhesion (Raucher *et al.*, 2000), promotes the production of actin filaments within the cell (Janmey *et al.*, 2018), and is capable of removing capping proteins like CapZ from actin filaments, to amplify growth (Schafer *et al.*, 1996; Toker, 1998). Furthermore, PIP_2_ is able to interact with several actin-binding proteins such as geloslin (Janmey and Stossel, 1987), cofilin (Yonezawa *et al.*, 1990) and Ezrin/Radixin/Moesin (ERM) proteins (Hirao *et al.*, 1996), with ERM proteins working to crosslink the PM and actin filaments (Arpin *et al.*, 2011). Several actin-binding proteins found in purified influenza virus (Shaw *et al.*, 2008) have known PIP_2_-interactions (Catimel *et al.*, 2008; Catimel *et al.*, 2009). The relationship between PIP_2_ and actin filaments further implicates PIP_2_ as having a role in additional cellular processes like chemotaxis and cell migration (Wu *et al.*, 2014). PIP_2_ also plays a role in allowing the adapter protein AP-2 to associate with clathrin-coated pits (Gaidarov and Keen, 1999). Any of these interactions could potentially be affected by CPC, both in the context of viral infection and normal cell function, and should be examined in future research.

A plethora of studies in different viruses show that phosphoinositides, and in many cases PIP_2_, interact with viral proteins through multiple basic residues, and that these interactions are crucial for the life cycles of multiple viruses (Ono *et al.*, 2004; Chukkapalli *et al.*, 2008; Chukkapalli and Ono, 2011; Altan-Bonnet and Balla, 2012; Johnson *et al.*, 2016; Yandrapalli *et al.*, 2016; GC *et al.*, 2017; Budicini *et al.*, 2018). More recently, phosphoinositide kinase inhibitors, which affect cellular PIP_2_ levels, have been shown to inhibit Zaire Ebola virus and SARS-CoV-2 (Kang *et al.*, 2020). Phosphoinositide kinase inhibitors also inhibit mast cell degranulation (Santos *et al.*, 2013). Targeting interactions between viral proteins and host cell phosphoinositides could be a novel therapeutic approach; CPC is already being used in a clinical trial against the SARS-CoV-2. Our study presents the effect of CPC on PIP_2_-binding proteins and illuminates a mechanism for its antiviral properties at relatively non-cytotoxic concentrations, paving the path for the future study of the effect of CPC in modulating PIP_2_ and PIP_2_-binding proteins and other membrane proteins that interact with the PIP_2_.

In future work, we aim to further deeply explore the CPC, PIP_2,_ and HA interactions in multiple cell types including polarized epithelial cells. Future virology research will be necessary to pinpoint the exact mechanism of action of CPC anti-influenza action within the viral lifecycle.

In conclusion, here we report that low-dose CPC acts as an effective anti-influenza agent in zebrafish. The underlying mechanism of action is likely related to the ability of CPC to disrupt HA cluster density, the number of HA molecules per cluster, and the nanoscale co-localization of HA with the PIP_2_ reporter PAmKate-PH, which we infer is due to CPC interference with PIP_2_. In fact, CPC disrupts multiple (MARCKS, PH, and HA) PIP_2_-dependent protein interactions and subcellular localizations. Super-resolution microscopy makes possible these mechanistic investigations. CPC also interferes with PIP_2_-dependent cellular functions (mast cell degranulation) and organismal interaction with influenza (as evidenced by CPC reduction of zebrafish mortality and viral burden). In summary, CPC targets the key eukaryotic signaling lipid PIP_2_ and reduces influenza morbidity by modulating host cell-viral protein interactions.

## Abbreviations

Ag: antigen
ATP: adenosine triphosphate
BSA: bovine serum albumin
BT: Tyrodes-bovine serum albumin
CMC: critical micelle concentration
CPC: cetylpyridinium chloride
CT: cytoplasmic tail
DC: duct of Cuvier
DNP: anti-dinitrophenyl
dpf: days post fertilization
dpi: days post infection
ER: endoplasmic reticulum
FPALM: fluorescence photoactivation localization microscopy
HA: hemagglutinin
hpf: hours post fertilization, hpi, hours post-infection
IAV: influenza type A virus
IgE: immunoglobulin E
IP_3_: inositol 1,4,5-triphosphate
LDH: lactate dehydrogenase
MARCKS: myristoylated alanine-rich C-kinase substrate
MDCK: Madin-Darby Canine Kidney
NIH-3T3: mouse embryo fibroblast cells, 3-day transfer, inoculum 3×10^5^ cells
PBS: phosphate buffered saline
PIP_2_: phosphatidylinositol 4,5-bisphosphate
PLCγ: phospholipase C gamma
PH: pleckstrin homology
PM: plasma membrane
RBL-2H3: rat basophilic leukemia cells, clone 2H3
ROI: region of interest
SEM: standard error of the mean
SARS-CoV-2: severe acute respiratory syndrome coronavirus 2
SOCE: store-operated Ca^2+^ entry.

## Acknowledgments

We thank Dr. Robert Wheeler, Siham Hattab, and Bailey Blair for use of and help with the ibidi heating system and confocal microscope; Dr. Joshua Kelley for guidance regarding image analysis; Dr. Juyoung Shim for helpful discussions, and Liz Saavedra Perez for the confirmation of TCID_50_ assay results, and Patricia Byard for administrative assistance.

This research was supported by the National Institutes of Health: National Institute of General Medical Sciences award numbers P20GM103423 (an Institutional Development Award) and 1R15GM139070 (PI: Hess), and R15GM116002 (PI: Hess), as well as the National Institute of Allergy and Infectious Diseases award number R15AI131202 (PI: King), and by the Maine Technological Asset Fund (MTAF 1106 and 2061, PI: Hess). University of Maine funding that supported this work includes Office of the Vice President for Research, the Maine Economic Improvement Fund, the University of Maine System Research Reinvestment Fund Grant Program Track 1 Rural Health and Wellbeing Grand Challenge (PI: Gosse), a Regular Faculty Research Grant (PI: Gosse), a UMaine Medicine Seed Grant (PI: Gosse), a Graduate Student Government Grant (PI: Obeng), a Charlie Slavin Research Grant, Frederick Radke Undergraduate Research Fellowships, Maine Top Scholar research supply funds, and the Center for Undergraduate Research.

## Supplemental Material

**Figure S1.**
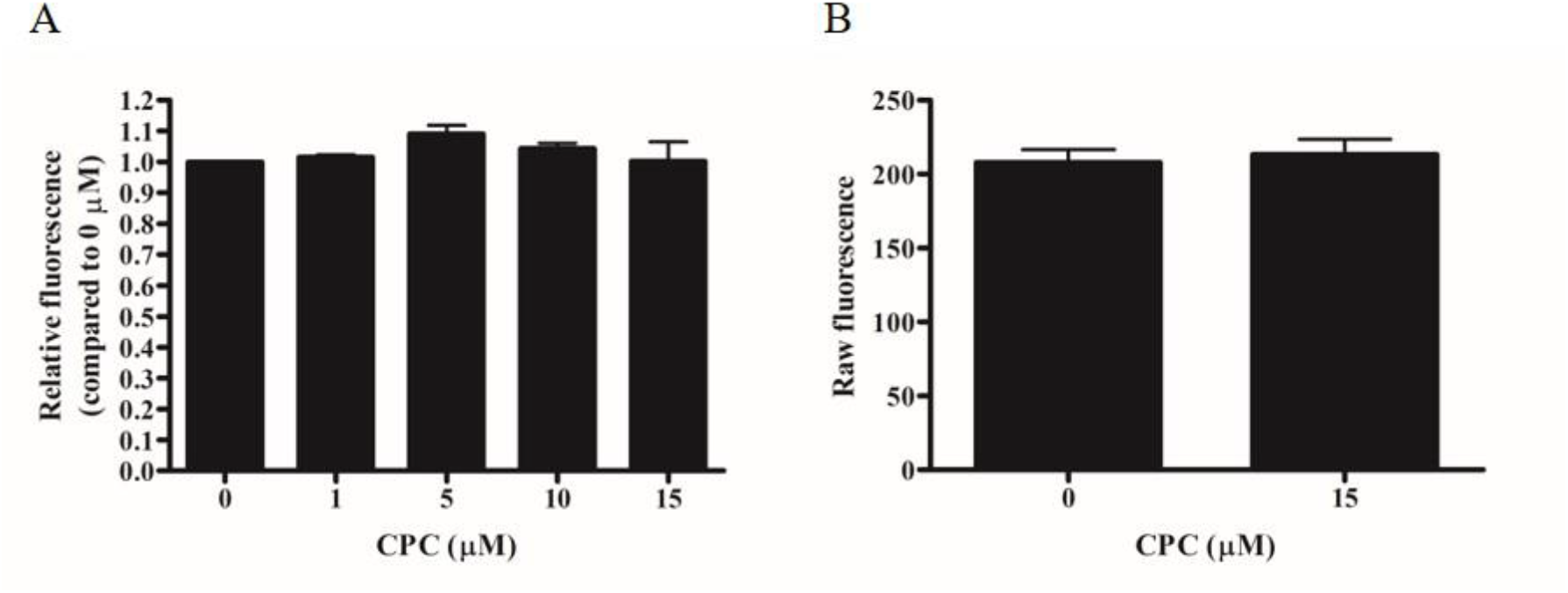
Effect of CPC on the ability of β-hexosaminidase to cleave fluorogenic substrate and effect of CPC on background fluorescence. **(A)** This is a control to test for CPC interference with the enzyme-substrate reaction between β-hexosaminidase and 4-methylumbelliferyl-N-acetyl-ß-D-glucosaminide (4-MU) in the assay used to quantify mast cell degranulation. **(B)** This is a control to test for CPC interference with background fluorescence in the degranulation assay. **Methods: (A)** Supernatant from a flask of degranulated cells, which consists of β-hexosaminidase but no cells, was combined in equal proportions with CPC at the doses indicated and incubated for 30 min at 37 °C. This way, CPC was present only during the enzyme-substrate step of the degranulation measurement. Triplicate 25-μL samples of each supernatant-CPC mixture were plated into a Grenier 96-well black-bottom plate containing 100 μL per well of 1.2 mM 4-MU in sodium acetate buffer (Hutchinson *et al.*, 2011). Tyrode’s-bovine serum albumin was used for background wells. The plate was incubated for 30 min at 37 °C, and 200 μL of glycine carbonate buffer was subsequently added to each well. Fluorescence was quantified using a Synergy2 microplate reader (Biotek). Background-subtracted fluorescence values were normalized to 0 μM CPC. **(B)** CPC at the doses indicated were plated in triplicate into background wells (no cells, no IgE, no Ag), and raw fluorescence values were measured in the microplate reader (Biotek). **Results: (A)** Values presented are means ± SEM for three independent experiments. No statistical significance was demonstrated by a one-way ANOVA followed by Tukey’s post-test (p < 0.05), compared to 0 μM CPC. CPC had no effect on β-hexosaminidase activity, indicating that the mast cell degranulation assay measuring β-hexosaminidase release is sound. **(B)** No statistical significance was demonstrated by a one-way ANOVA followed by Tukey’s post-test (p < 0.05), compared to 0 μM CPC.

**Figure S2.**
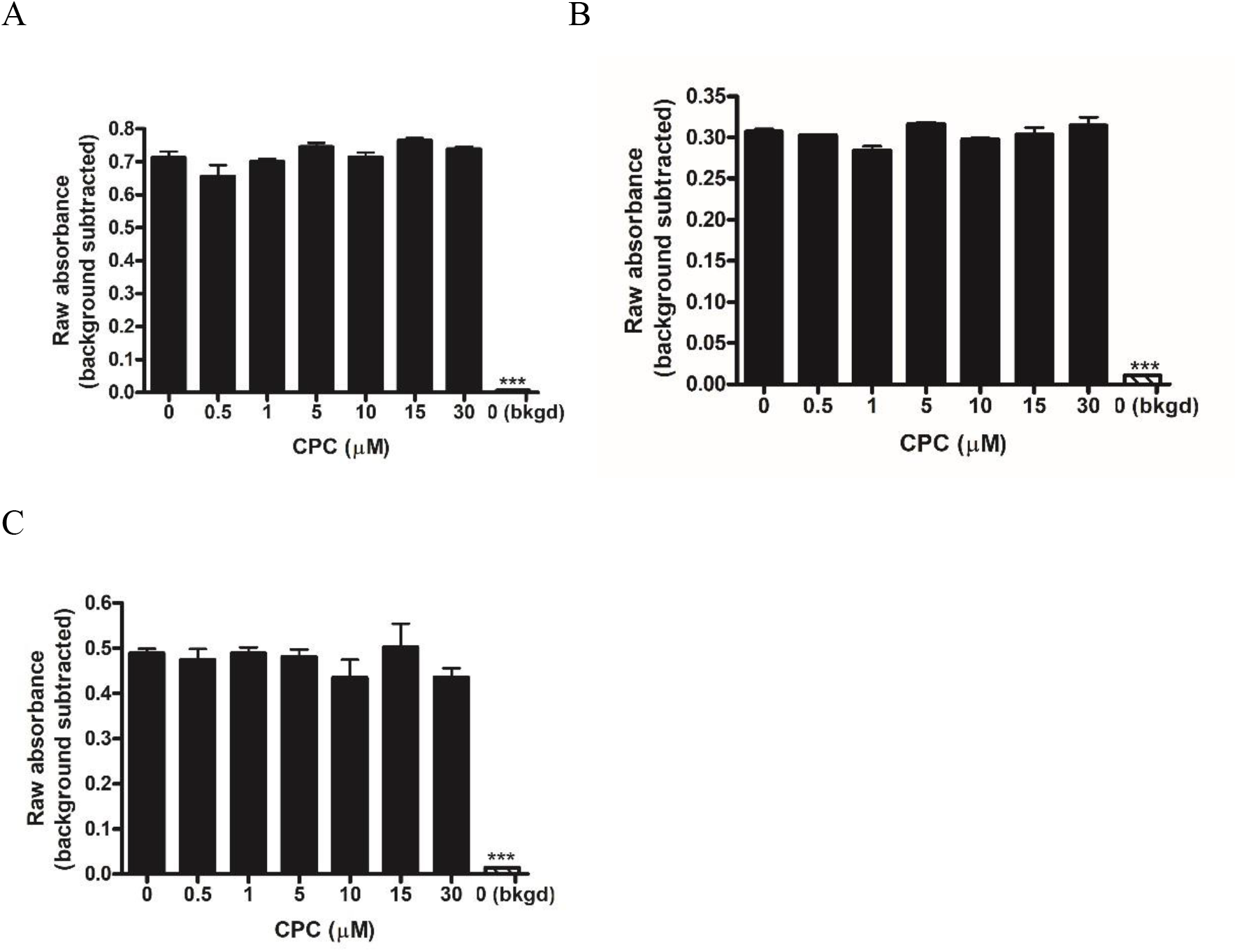
No-cell control to assess whether CPC interferes with the LDH enzymatic reaction that is used as a cytotoxicity readout. CPC at the doses indicated was plated with **(A)** 0.0003125 U/mL or **(B)** and **(C)** 0.000625 U/mL of purified LDH enzyme (Cayman chemical company, Ann Arbor, MI, USA). In a 96-well plate, 50 μL of a 2X solution of CPC (in BT) was plated with 50 μL of 2X LDH solution. Only BT (no LDH or CPC) was plated into the background wells. Following a 1-h incubation at 37 °C, the LDH cytotoxicity kit (Roche) was used per manufacturer’s instructions to quantify LDH enzyme activity. Absorbance values at 490 nm and background absorbance at 630 nm were recorded, and the background values were subtracted from the 490 nm values. Values represent the averages of triplicate samples for individual experiments. Error bars represent SEM. Significance determined using a Tukey’s post-test and one-way ANOVA. *** p < 0.0001.

**Figure S3.**
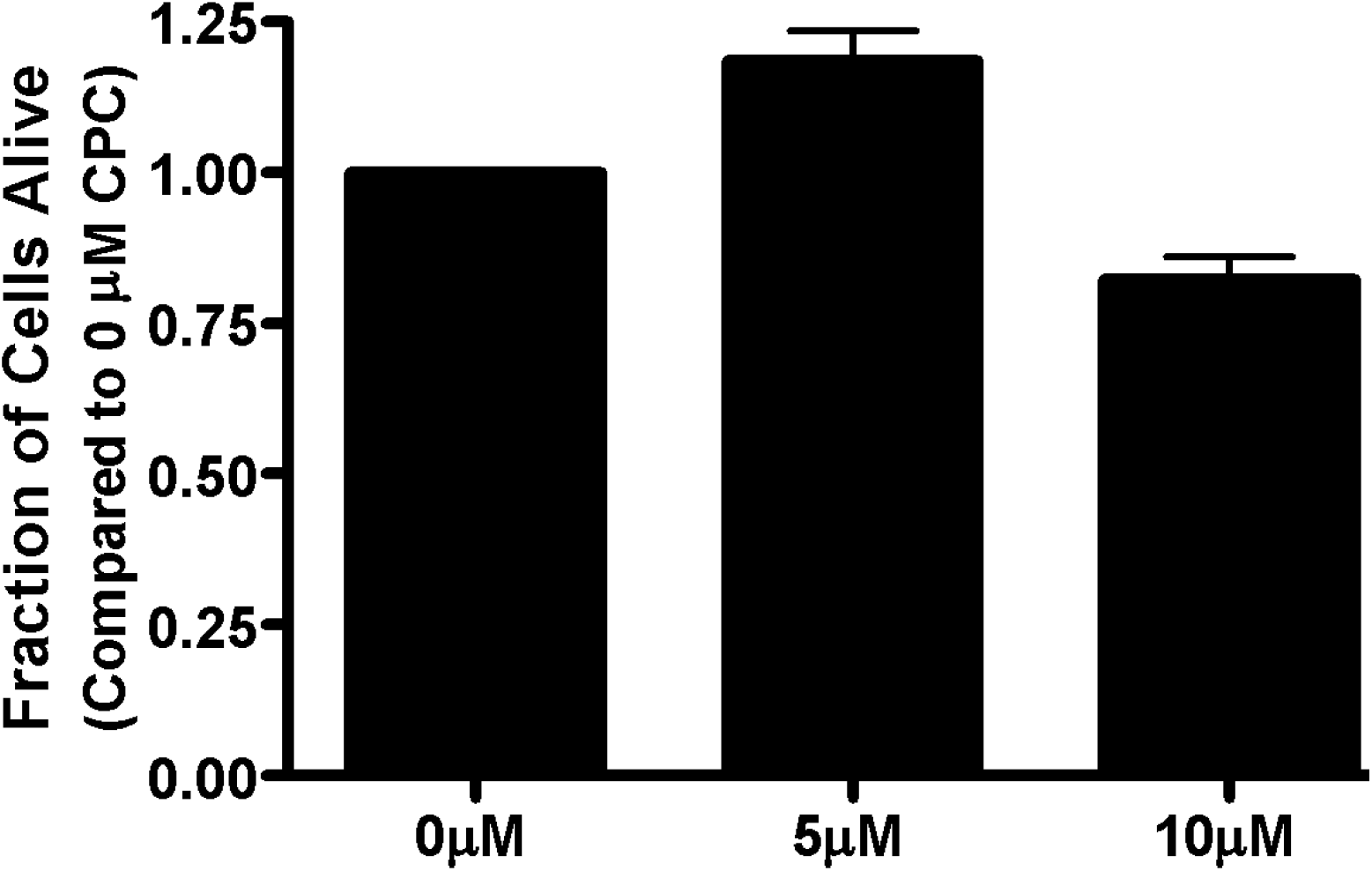
Cytotoxic effects of CPC on NIH-3T3 cells. CPC cytotoxicity was tested in NIH-3T3 cells to add context to microscopy data utilizing this cell line. CPC was prepared as in Methods. NIH-3T3 cells were cultured as in (Weatherly *et al.*, 2016). Cells were plated at 200,000 cells per well into each of 3 wells of a 6 well plate (Greiner) and grown overnight at 37 °C/5% CO_2_. Next day, cells were exposed to micromolar doses of CPC (0 µM, 5 µM, and 10 µM) and incubated for 1 hour at 37 °C/5% CO_2_. Cell viability was assessed using a trypan blue exclusion assay (as in section 2.1.4). Values presented are means ± SEM of three independent experiments. Analysis by the nonparametric Kruskal-Wallis test followed by Dunn’s post test found no statistical significance of either CPC dose compared to control. CPC (up to 10 µM, 1 hr) exposure does not cause cytotoxicity of NIH-3T3 cells.

**Figure S4.**
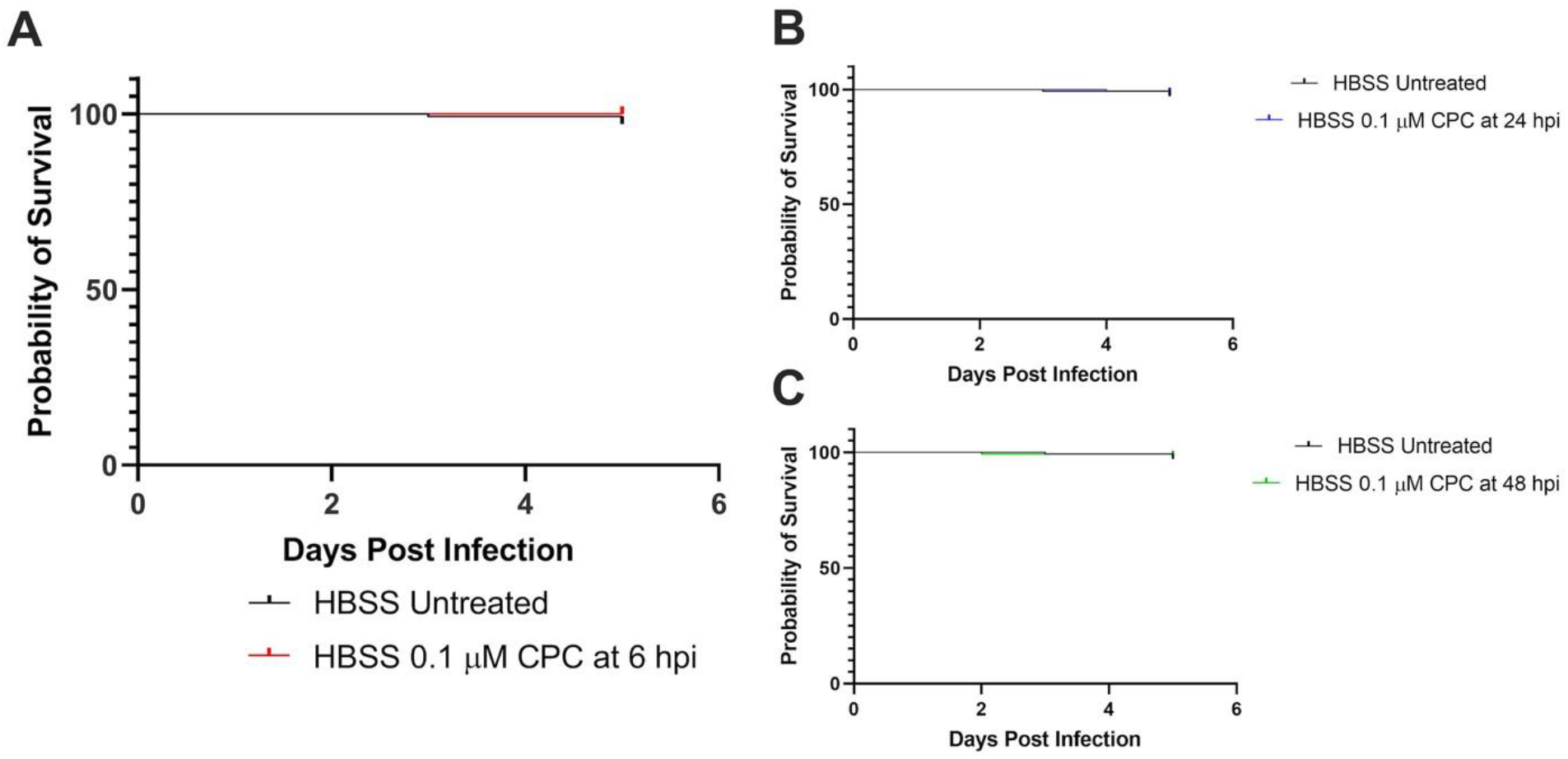
No significant mortality for HBSS-injected embryos. **(A)** Survival curve between 0.1 µM CPC treated and untreated, HBSS-injected zebrafish with a one-hour CPC treatment administered at 6 hpi. **(B)** Survival curve between 0.1 μM CPC treated and untreated HBSS injected zebrafish with a one-hour CPC treatment administered at 24 hpi. **(C)** Survival curve between 0.1 μM CPC treated and untreated HBSS injected zebrafish with a one-hour CPC treatment administered at 48 hpi. By 5 dpf the average survival of CPC treated zebrafish and untreated plates showed no significance in mortality within a 2.5% range. All graphs represent an n of 3 with ∼150 zebrafish compared.

**Figure S5.**
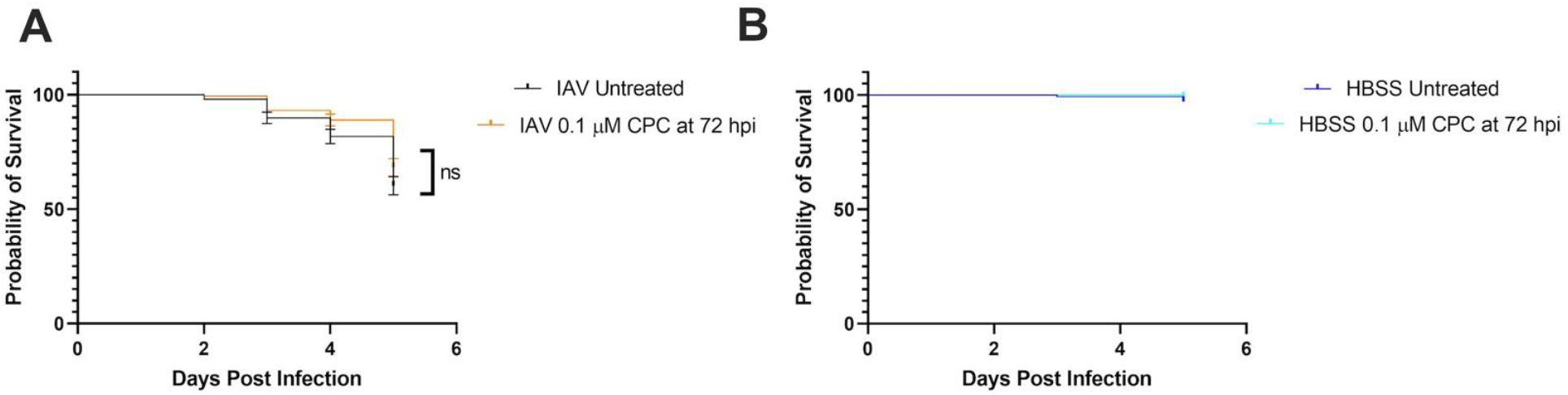
No significant mortality between HBSS and IAV at 72 hpi. **(A)** Survival curve between 0.1 μM CPC treated and untreated IAV infected zebrafish with CPC treatment administered at 72 hpi. There was no significant change in mortality between untreated and treated IAV infected groups. **(B)** Survival curve between 0.1 μM CPC treated and untreated HBSS injected zebrafish with CPC treatment administered at 72 hpi. There was no significant change in mortality between untreated and treated HBSS groups.

